# Automating data analysis for hydrogen/deuterium exchange mass spectrometry using data-independent acquisition methodology

**DOI:** 10.1101/2023.08.25.554852

**Authors:** Frantisek Filandr, Vladimir Sarpe, Shaunak Raval, D. Alex Crowder, Morgan F. Khan, Pauline Douglas, Stephen Coales, Rosa Viner, Aleem Syed, John A. Tainer, Susan P. Lees-Miller, David C. Schriemer

## Abstract

We developed a hydrogen/deuterium exchange workflow coupled to tandem mass spectrometry (HX-MS^2^) that supports the acquisition of peptide fragment ions alongside their peptide precursors. The approach enables true auto-validation of HX data by mining a rich set of deuterated fragments, generated by collisional-induced dissociation (CID), to simultaneously confirm the peptide ID and authenticate MS^1^-based deuteration calculations. The high redundancy provided by the fragments supports a confidence assessment of deuterium calculations using a combinatorial strategy. The approach requires data-independent acquisition (DIA) methods that are available on most MS platforms, making the switch to HX-MS^2^ straightforward. Importantly, we find that HX-DIA enables a proteomics-grade approach and wide-spread applications. Considerable time is saved through auto-validation and complex samples can now be characterized and at higher throughput. We illustrate these advantages in a drug binding analysis of the ultra-large protein kinase DNA-PKcs, isolated directly from mammalian cells.

## Introduction

Hydrogen/deuterium exchange coupled to mass spectrometry (HX-MS) is a labelling method based on the exchange between protein backbone amide hydrogens and deuterated labelling buffers. HX-MS is used to obtain information about higher-order protein structure and dynamics, as the rate of amide hydrogen exchange is influenced by local structure and solvent accessibility. For example, it can help investigate protein folding mechanisms^1^, discover ligand binding sites and highlight allosteric effects of binding^2^. In the biotherapeutics industry it is particularly useful for epitope mapping^3^. The use of native solution conditions during labelling and the low sample requirements of the method makes HX-MS an appealing technology for structure-function analysis. Even complex multi-protein systems can be interrogated, if the system produces a suitable number of peptides in the sample workup process. The method has been reviewed extensively in recent years^4–12^. The standardization of experimental and data reporting protocols have improved the accessibility of the technology, creating the reliable biophysical technique that we know today^13^.

However, while HX-MS enjoys frequent use in protein analysis, it is generally restricted from applications involving high-throughput characterizations, or analyses involving protein states much larger than 150kDa of unique sequence^6^. This restriction is no longer due to limitations in the analytical systems. Advancements in ion mobility can reduce spectral complexity^14,15^ and new methods even support nanoHX on ultra-large complexes^16^. Robotic technologies remove much of the complexity of data collection^17–19^ and even sub-zero chromatography can extend the window of elution time and minimize the problem of deuterium back exchange^20^.

The remaining bottleneck is data analysis. Complex isotopic profiles are generated by the technique and even with the improvements mentioned above, spectral overlap still occurs and deuteration values can be miscalculated. These worsen as sample complexity increases. We seek good quality mass spectra with clean peptide isotopic envelopes when collecting deuteration data. To date, the field manually inspects the raw data to validate spectral selections and discard peptide signals that are compromised in any way. Retention time is used to confirm the identity of a peptide, supported by the accurate mass of a deuterium-shifted signal and signal “quality” is visually assessed, relying upon years of experience in spectral assessment. But as the size of the protein system grows, or the number of states to screen increases, manual curation becomes impractical and such an approach has always been prone to human error. A solution is needed that would remove the burden of manual data validation entirely, while also tolerating more convoluted spectra arising from complex mixtures, even whole cell lysates. Unfortunately, all major data analysis engines only focus on supporting manual review and post-validation activities such as data visualization and statistical analysis^21–27^.

One possible strategy for automated analysis involves leveraging peptide fragmentation and the MS^2^ domain. The acquisition of fragments in an HX-MS^2^ experiment can corroborate the identity of a peptide and generate abundant data to confirm the deuteration level of the precursor peptide. The approach relies upon deuterium scrambling, which is ubiquitous under normal ion transmission conditions. We had previously demonstrated the potential of such an approach^28^. However, we lacked an efficient way to collect and mine the data in a comprehensive and platform-independent manner. In this work, we demonstrate how data independent acquisition (DIA) can be used for HX-MS^2^ experiments as a method to obtain deuteration data from both MS^1^ and MS^2^ domains simultaneously (**Fig. 1**). We adapt computer vision algorithms that allow us to automatically validate the selection of peptides for deuteration analysis and even rescue overlapped signals. The resulting analyses are comparable to expert curated datasets, while offering objectively validated data and a clear measure of reliability for each peptide datapoint. Export-ready figures in the form of uptake plots and differential Woods plots are automatically generated at the end of data processing. We demonstrate that highly complex samples drawn directly from cell lysates can be accurately interrogated in minutes, rather than days or weeks.

**Figure 1.**
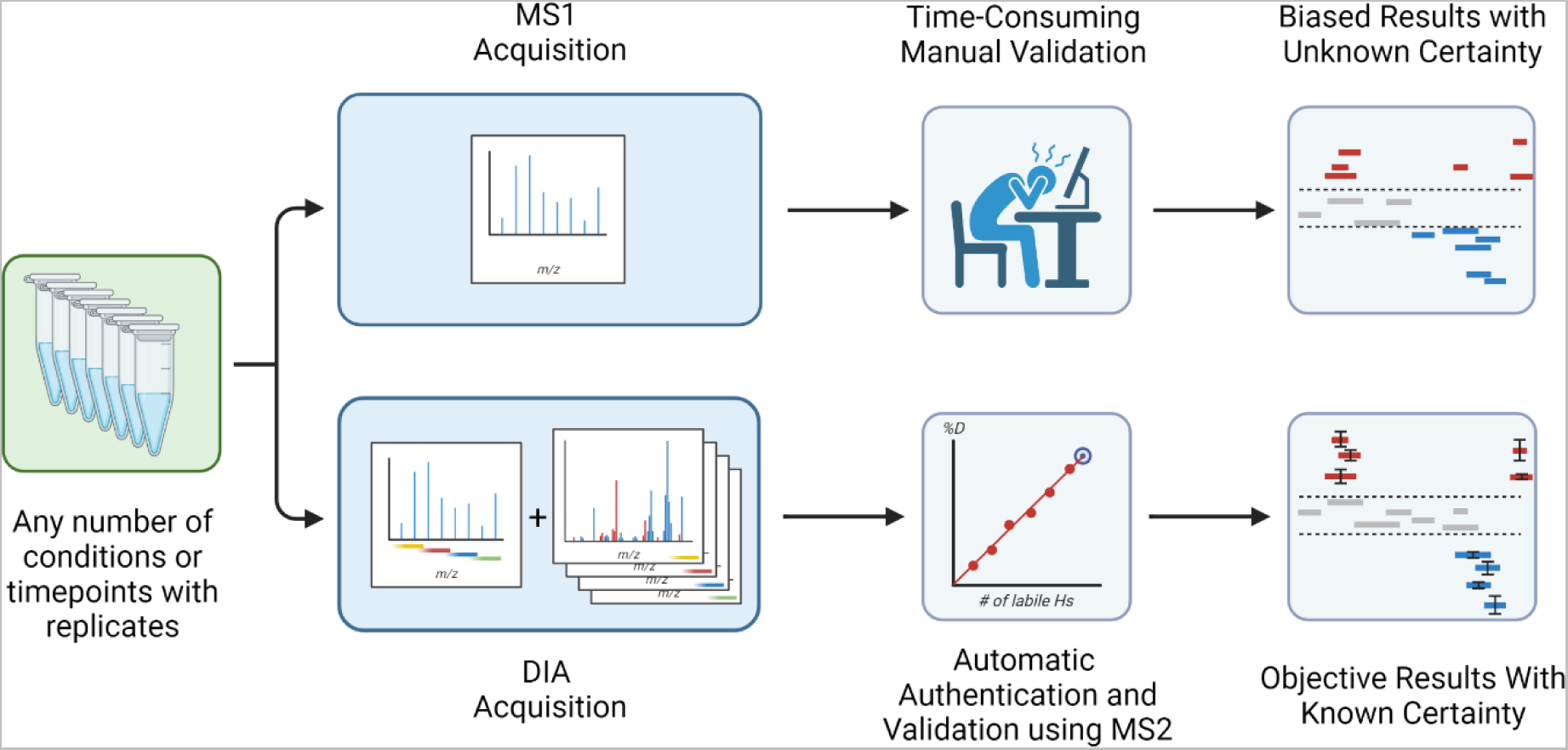
DIA acquisition enables automation. DIA data acquisition uses deuterium-scrambled CID fragments as surrogates that confirm the identity and the deuteration value of any given peptide. It supports the implementation of an automation approach to generate Woods plots or deuterium uptake curves without any user input required, while also offering data reliability for each peptide based on fragment statistics.

## Experimental Setup

### HX-MS^2^ of phosphorylase B

#### D_2_O labelling

Phosphorylase B (Sigma-Aldrich, P6635) was resuspended in HEPES buffer (25 mM, pH 7.4, 150 mM NaCl) to a concentration of 10 µM and diluted with deuterated HEPES buffer (25 mM, pD 7.4, 150 mM NaCl) to create a 50% D_2_O labelling mixture. Labeling was conducted over multiple timepoints from 15 seconds to 1 hour, and aliquots were quenched 1:1 (v/v) with 250 mM glycine buffer (pH 2.3) containing 0.5 µg/µl nepenthesin II, resulting in a 1:1 protein:protease ratio (w/w). Samples were digested at 8°C for 2.5 minutes and then flash frozen in liquid nitrogen. Samples were prepared in triplicate for each timepoint and thawed immediately prior to HX-MS^2^ analysis. Samples for sequence mapping based on data dependent acquisition (DDA) were processed in the same way, except D_2_O was replaced with H_2_O in the labelling phase.

#### Data Collection

Data were acquired on a Thermo Scientific Q Exactive^TM^ Plus mass spectrometer connected to a LEAP PAL HDX autosampler and a Thermo Scientific UltiMate^TM^ 3000 LC system. All samples were prepared manually and injected into the cold compartment of the autosampler set to 4°C. The injected peptides were trapped on a Luna 5µm C18(2) 100Å Micro Trap (20 x 0.50 mm, Phenomenex) and separated on a Luna Omega 3 µm Polar C18 100 Å LC Column (50 x 0.3 mm, Phenomenex) using a standard 5% to 45% solvent B gradient (10 min. gradient). Solvent A was 0.4% FA in H_2_O and solvent B was 0.4% FA in 80% ACN. The flowrate was 15 µl/min for separation and 70 µl/min for loading and desalting. Data were collected using a standard HESI source. The spray voltage was set to 3500V, sheath gas flow rate to 20, auxiliary gas flow rate to 7, and sweep gas flow rate to 0. The capillary temperature was set to 250°C, the S-lens RF level to 65 and the auxiliary gas heater temperature to 80°C. Approximate chromatographic peak widths were 13 sec (FWHM) with this configuration.

DDA mapping runs were performed according to previously published optimized settings^29^ with the top 12 ions selected for fragmentation. MS resolution was set to 70 000 with an AGC target of 5e5 and a 120 msec maximum trap fill time. The mass range was set to m/z 300-1040. MS^2^ scans were collected at a resolution of 35 000 with an AGC target of 5e5 and a 120 msec maximum trap fill time. The isolation window was set to m/z 2.5 with a collision energy of 28 NCE. Total cycle time was approximately 1.8 seconds. Dynamic exclusion was set to 9 seconds to allow for approximately two MS^2^ scans of each chromatographic feature.

DIA acquisitions of deuterated samples consisted of a single MS^1^ scan of m/z 300-1040 followed by a set of 16 fragmentation bins with a width of m/z 50 and an overlap of m/z 4 per bin edge, covering the whole mass range. Full MS scan resolution was set to 70 000 with an AGC target of 1e6 and a 50 msec maximum trap fill time. DIA scans were set to a resolution of 35 000 with an AGC target of 2e5 and a 120 msec maximum trap fill time. The fixed first mass was set to m/z 200 and the collision energy to 28 NCE.

### HX-MS^2^ of Polϴ ± novobiocin

#### D_2_O labelling

For differential HX-MS^2^ analysis, the ATPase domain of Polθ was produced in baculovirus-infected insect cells as previously described^30^. A 4 µM solution in HEPES buffer (25 mM, pH 7.4, 250 mM NaCl) was mixed 1:1 with 4 mM novobiocin (prepared in the same HEPES buffer with 2% DMSO) and incubated for 30 min. For each sample, 5 µL of the pre-incubated mixture was combined with 5 µL of D_2_O-based HEPES buffer (25 mM, pD 7.4, 250 mM NaCl) to initiate deuterium labelling at room temperature. After 2 min of labelling, the reactions were quenched 1:1 (v/v) with quench/digestion buffer (500 mM glycine pH 2.3, 6 M urea) containing 0.6 µg/µL nepenthesin II digestion enzyme. Samples were digested at 8°C for 2 minutes and then flash frozen in liquid nitrogen. Control samples without novobiocin were prepared with matched DMSO concentrations and processed as above, but with a quench/digestion buffer containing just 0.2 µg/µL nepenthesin II. All samples and controls were prepared in triplicate.

#### Data Collection

Data were acquired on a Sciex TTOF 6600 instrument with an Optiflow Nano ESI Source, integrated with a Sciex Ekspert nanoLC 425 and a Trajan PAL HDX autosampler. Samples were manually injected into the cold compartment of the autosampler (set to 4°C), outfitted with an Acclaim™ PepMap™ 100 C18 HPLC trap column for desalting (0.1 mm diameter, 5 µm particle size, 100 Å pore size, 20 mm length) at 10 µL/min mobile phase A for 3 min. Concentrated sample was eluted and separated on a nanoEase M/Z Peptide CSH C18 Column (75 µm diameter, 1.7 µm particle size, 130 Å pore size, 150 mm length), connected directly to the ion source, using a linear 10-minute gradient from 5%-35% mobile phase B at 250 nl/min. Source settings were as follows: GS1 = 7, GS2 = 0, CUR = 25, TEM = 0, ISVF = 3800. Approximate chromatographic peak widths were 6 sec (FWHM).

DDA mapping runs were performed with the top 10 ions selected for fragmentation in “high sensitivity” mode. The MS^1^ mass range was set to 400-850 m/z with a 150 ms accumulation time. The MS^2^ scan range was set to 350-1100 m/z with an accumulation time of 120 ms and scans were collected with a dynamic accumulation, dynamic collision energy and dynamic background subtraction functions enabled. The total cycle time was approximately 1.4 seconds. Dynamic exclusion was set to 20 seconds, allowing only a single MS^2^ acquisition of all chromatographic features.

HX-MS^2^ runs were performed with 16 variable DIA windows with m/z 3 overlap in “high sensitivity” mode. The MS^1^ mass range was set to 400-850 m/z with a 200 ms accumulation time. DIA scan range was set to m/z 350-1100 with an accumulation time of 100 ms and scans were collected with a rolling collision energy setting, optimizing collision energy for each DIA window. The total cycle time was approximately 1.85 seconds.

### HX-MS^2^ of DNA-PKcs ± AZD7648

#### Coupling of anti-GFP nanobodies to magnetic beads

0.5 mg of Dynabeads™ (MyOne™ Streptavidin T1) were incubated with 10 µg of biotinylated Alpaca anti-GFP nanobody (ChromoTek, GTB-250) in a 100 µL incubation volume (PBS, pH 7.4, 0.1% Triton X-100) for 4 hours at 4 °C. After conjugation, beads were washed three times with 100 µL of incubation buffer to wash away unbound nanobody.

#### Expression and affinity enrichment of EGFP-DNA-PKcs construct

Human EGFP-DNA-PKcs construct stably expressed in DNA-PKcs null V3 CHO cells was a kind gift from Dr. Kathy Meek (Michigan State University). The detailed preparation of V3 transfectant is described elsewhere^31^. The cells were cultured in 10 cm plates in α-MEM supplemented with 10% fetal bovine serum, 100 U/mL penicillin and streptomycin, 10 µg/µL cipromycin, and 10 µg/µL blasticidin. Cells were harvested by trypsinization and washed twice with 10 mL of PBS. Cell pellets were frozen at −80 °C until protein extraction. Frozen cell pellets were suspended in 1 mL of lysis buffer (50 mM Tris-HCl, 150 mM NaCl, 1 mM EDTA, 0.5% NP-40) with cOmplete™ protease inhibitor cocktail (Roche), phosphatase inhibitor cocktail (Roche), and universal nuclease (Pierce). The lysate was incubated on a nutator for 30 minutes at 4 °C followed by sonication with 3 × 5 seconds burst on ice. Lysate was centrifuged at 14,000g, for 15 minutes at 4 °C, and protein concentration in clarified cell lysate was determined using BCA protein assay (Pierce). The protein concentration was adjusted to 2 mg/mL and 2 mg aliquots were flash frozen at −80 °C until pulldown. 80 µg of anti-GFP nanobody conjugated Dynabeads™ (approximately 45 nL) were incubated with the cell lysate (4 mg of total protein content) prepared from the EGFP-DNA-PKcs expressing V3 CHO cells for 90 minutes at 4 °C. The beads were isolated and washed with 250 µL of PBS (pH 7.4) on a magnetic nano-isolator device described elsewhere. Beads were removed from the nano-isolator and collected in 8 µL HEPES buffer (10 mM, pH 7.4).

#### D_2_O labelling

Prior to deuterium labelling, 10 µg of isolated beads (approximately 6 nL) were mixed in 4.5 µL of equilibration buffer (10 mM HEPES, pH 7.4) ± AZD7648 (Selleckchem cat # S8843, 1 µM) for 10 minutes. The deuterium labelling was carried out for 5 minutes by addition of 4.5 µL of D_2_O labelling buffer (10 mM HEPES, pD 7.4). Labelling was quenched with addition of 1 µL of digestion buffer (500 mM Glycine-HCl, pH 2.3) containing 0.6 µg/µL of Nepenthesin II digestion enzyme. Digestion was carried out for 90 seconds at 10 °C. Following digestion, the beads were magnetized, and the digest was collected and flash frozen. To determine the protein composition in the pulldown, 10 µg of isolated beads were incubated with MS-grade trypsin (15 ng, 50 mM AMBIC, pH 8.0) for an overnight on-bead digestion at 37 °C. The next morning, digest was quenched with the addition of 1 µL of 20% FA. The beads were magnetized, and the digest was collected in a sampling vial for MS analysis.

#### Data Collection

Data were collected on a prototype nanoHX ion source^16^ coupled with an Orbitrap Eclipse, outfitted with a Vanquish Neo loading pump and a Vanquish Neo gradient pump. Samples were manually injected into the chilled nanoHX source (held at 4°C), which contained a PepMap™ Neo C18, 5 µm 300 µm x 5 mm trap cartridge (Thermo Fisher Scientific, P.N: 174500) and a PepMap™ Neo 2 µm C18, 75 µm x 150 mm analytical separation column (Thermo Fisher Scientific, P.N: DNV75150PN). Peptides were loaded and washed at 50 µl/min (0.4% formic acid) for 1 minute. The concentrated sample was eluted using a linear 25-minute gradient from 0%-40% mobile phase B at 300 nl/min. The spray voltage was nominally set to 1700V with the time-dependent feature enabled. The RF lens was set to 30 and the ion transfer tube to 270°C. Approximate chromatographic peak widths were 6 sec (FWHM).

DDA mapping runs were performed in OT/OT mode with the MS resolution set to 60 000 and a mass range of m/z 375-1000. MS^2^ scans were collected at a resolution of 15 000 with isolation window set to m/z 1.6 with a collision energy of 30 NCE. Total cycle time was 1.5 seconds. Dynamic exclusion was set to 30 seconds after 2 occurrences within 15 seconds, to allow for approximately two MS^2^ scans of each chromatographic feature. The AGC target was set to standard and maximum injection time to auto.

DIA acquisitions of deuterated samples consisted of a single MS^1^ scan of m/z 350-1000 with a resolution of 60 000 followed by a set of 26 fragmentation bins with a width of m/z 25 and an overlap of m/z 4, covering the whole mass range with a resolution of 30 000. The fixed first mass in DIA was set to m/z 250 and the collision energy to 30 NCE. The AGC target was set to custom with a maximum injection time of 54 msec.

### Software Design and Availability

AutoHX functionality was built within the Mass Spec Studio 2.0 framework for integrative structural biology^32^. The software was written in C#, leveraging an extensive plugin-style repository of reusable content for rapid development of mass spectrometry applications. MSTools was used for workflow management^33^.

## Results and Discussion

### The properties of CID-generated HX-MS^2^ data

The fragmentation of deuterated peptides is accompanied by gas-phase scrambling of deuterium in a manner that is dependent upon the energetics of ion transmission and the mode of fragmentation used^34,35^. Scrambling is essentially complete using regular ion transmission settings as considerable ion activation occurs during desolvation and ion focusing^36^. Only when transmission conditions are detuned and used in conjunction with electron-mediated fragmentation modes can scrambling be reduced, as conventional CID fragmentation also contributes thermal energy and causes scrambling^37^.

Scrambling engages all sources of labile hydrogens in a peptide, and a full atom accounting reveals a linear relationship between the deuteration of a fragment and the number of its labile hydrogen sites (**Fig. 2**). The fragment deuteration model for the given deuterium-labeled peptide highlights a typical fit. This linear model intersects the origin and, in the absence of any spectral overlap in the MS^1^ domain, passes through the deuteration value of the precursor peptide. Selecting a single fragment is therefore sufficient to replace the precursor as an accurate and precise measure of deuteration, when scaled for size^28^. The longer and more intense fragment ions tend to be more sensitive measures of deuteration than smaller and less intense ones, but essentially all sequence ions can be used as surrogates for deuteration measurements, individually or combined. Only fragments that undergo neutral loss appear to deviate from the linear model, likely due to a kinetic isotope effect^28^.

**Figure 2.**
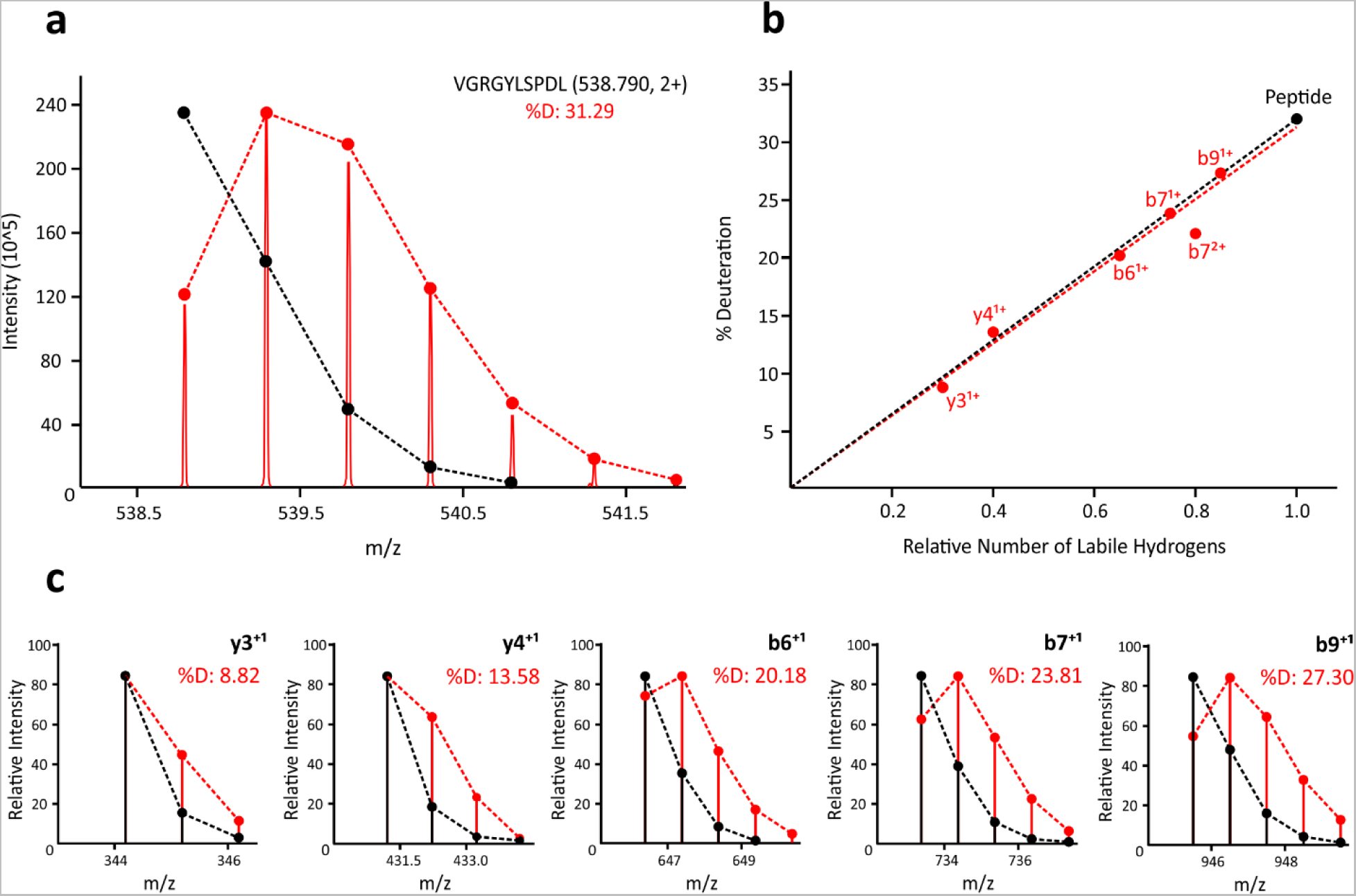
Effect of hydrogen/deuterium scrambling on fragment deuteration values for a given peptide. (**a**) MS^1^ spectrum of deuterated VGRGYLSPDL (2+ ion), showing native (black) and deuterium-expanded (red) isotopic envelopes. **(b)** Corresponding fragment deuteration model, where red dots represent fragment ion deuteration values and the black dot represents the precursor peptide deuteration. **(c)** Select fragment isotopic distributions supporting the model, showing native (black) and deuterium-expanded (red) isotopic envelopes.

### Building a DIA-based HX-MS^2^ workflow

Previous illustrations of fragment deuteration surrogacy used targeted MS^2^ acquisitions and data-dependent acquisition (DDA) experiments, together with limited deuteration experiments (*e.g.*, 20% D_2_O labelling)^28^. Reduced deuteration ensured that ions could be sampled within a conventional small ion transmission window (*e.g.,* m/z 2). A data-independent acquisition (DIA) experiment, with its wider transmission windows, should allow acquisition of fragment data for routine deuterium labelling experiments as we have previously suggested^38^. We refined our original HX-MS^2^ concept to implement this strategy.

We applied a standard DIA method design with one exception. To minimize spectral complexity in the MS^2^ spectra yet promote good sampling of chromatographic features, we restricted the mass range slightly and designed DIA ion transmission windows to be as small as possible for a given platform. The faster scanning the instrument, the smaller the windows can be made. However, unlike standard DIA methods, we use larger overlaps between successive windows to ensure that strongly deuterated peptides have at least one window where the peptide isotopic distribution is not truncated by the edge of a transmission window (**Fig. 3**), as such a truncation would result in distortion of resulting fragment isotopic envelopes due to missing isotopologues, and cause errors in the deuteration readout. A window size of m/z 4 was found to be sufficient and did not significantly compromise cycle times on either the TOF or Orbitrap platforms used in this study.

**Figure 3:**
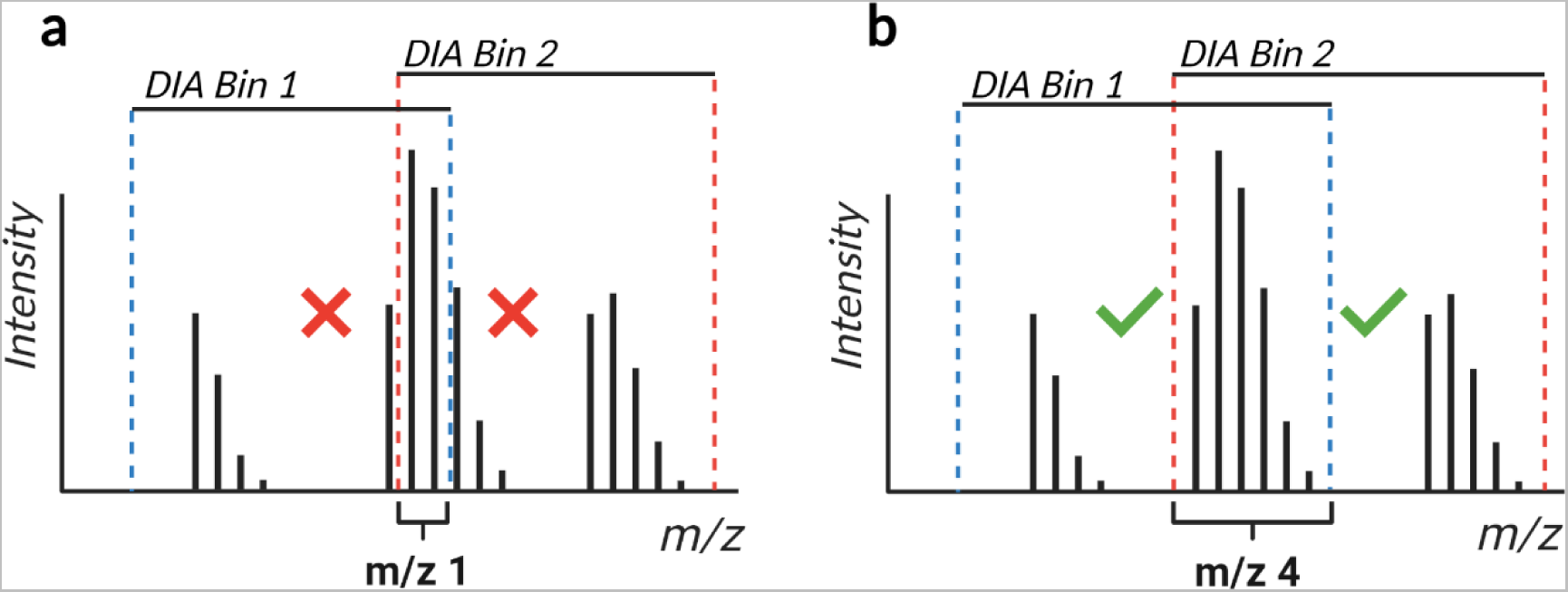
DIA window overlap overview. The standard m/z 1 DIA window overlap used in proteomics is insufficient if the goal is to obtain full and intact isotopic envelopes of fragment ions. If any of the isotopologues of the precursor peptide ion are cut out of the DIA bin, the resulting fragment isotopic envelopes will be distorted. An overlap of m/z 4 was found to be effective for a typical HX-MS experiment with most 2+ and 3+ charged peptides.

We developed AutoHX, a new software app in the Mass Spec Studio, to mine HX data in two dimensions. The software automatically selects the ideal, non-truncating DIA window for a given peptide and calculates deuteration values for the precursor peptide from the MS^1^ data and for all fragments from the MS^2^ data. The app currently requires a peptide library. This library is obtained from DDA runs that we collect at the beginning of an HX experiment, using matched but undeuterated control digests. These DDA runs can be searched with any standard peptide identification search engine. We revised HX-PIPE, our HX-tailored search engine^39^, to generate the library. HX-PIPE finds unambiguous peptide assignments and then formats a library for direct use in AutoHX. It has the option of generating a specific set of transitions from the peptide library, or deferring transition selection to AutoHX. We have found the latter to be more practical when fragmentation conditions vary slightly between the DDA and DIA runs (not shown).

Extracted ion chromatograms are then generated for each peptide in the library, and a window of integration is defined to produce averaged MS^1^ and MS^2^ spectra for deuteration analysis. A set of filters is applied to parse low quality signals from the dataset and then a RANSAC-based spectral analyzer is applied that selects the best set of isotopologues for all peptides and their fragments (**Fig. S1**). This spectral analyzer selects peaks based on a chosen deuteration model. EX2 is the current default but a more a complex EX1 model is also enabled. To determine the increase in peptide redundancy we obtain from a DIA-based deuteration measurement, we collected a triplicate, 6-timepoint kinetics analysis of phosphorylase B (a 97 kDa protein) in HX-DIA mode (**Fig. 4**). Analysis of the MS^1^ space generated 380 consistently useable peptide signals across all timepoints and replicates, whereas the fragment space generated 3269 useable fragment signals. Only peptides with 3 or more detectable quality fragments were used in this calculation. Thus, even though these data were collected on an older model instrument (QExactive Plus), adding the fragment dimension increases the redundancy in sequence coverage dramatically from 4.6 (MS^1^ only) to 49.7 (MS^1^ plus MS^2^). The value of this extra redundancy translates into better measurements. For example, calculating a deuterium value when including the fragment space generates a 41% improvement in measurement precision.

**Figure 4.**
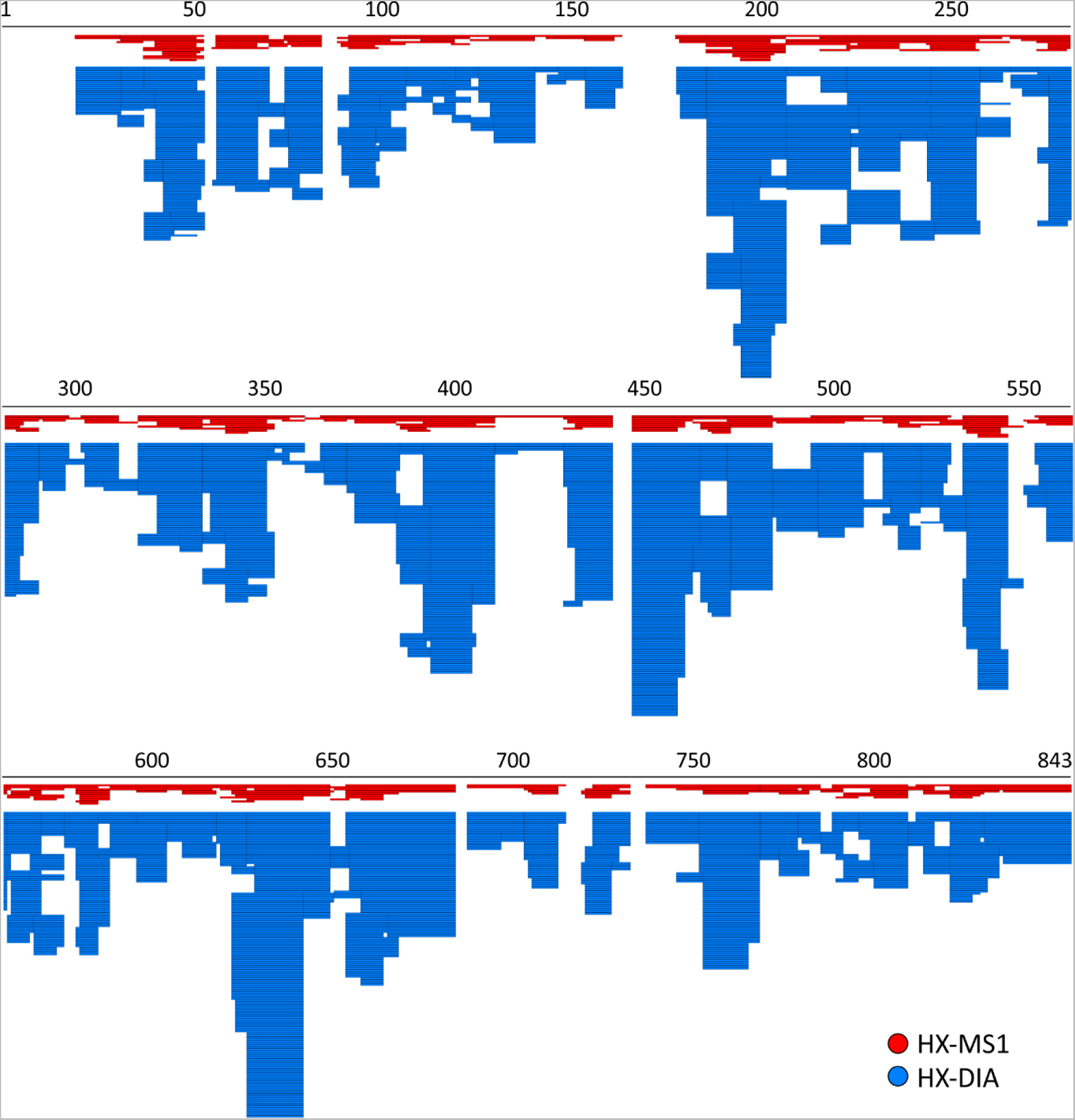
Deuteration map resulting from an HX-DIA kinetics analysis of phosphorylase B, showing the large increase in the redundancy of coverage obtained by including the fragment dimension in the analysis.

### Automated data authentication – kinetics

We noticed that there are instances where the MS^1^ measurement generates a more precise peptide deuteration measurement, and instances where a single fragment or even a combination of fragments generates a better measure. To automatically generate kinetic curves using the best of the underlying data in terms of accuracy and precision, we developed a method where all possible combinations of MS^1^ and MS^2^ data for a given peptide are created, producing a normal distribution of deuteration values (**Fig. 5**). The distribution is sampled and the combinations closest to the mean are mined for the one that generates the most precise deuteration value across the replicates. This combination is chosen to represent the peptide and is used in developing the kinetics curve. The final combination may differ between timepoints because the optimization step is done for every timepoint, to ensure that the cleanest signal is obtained.

**Figure 5:**
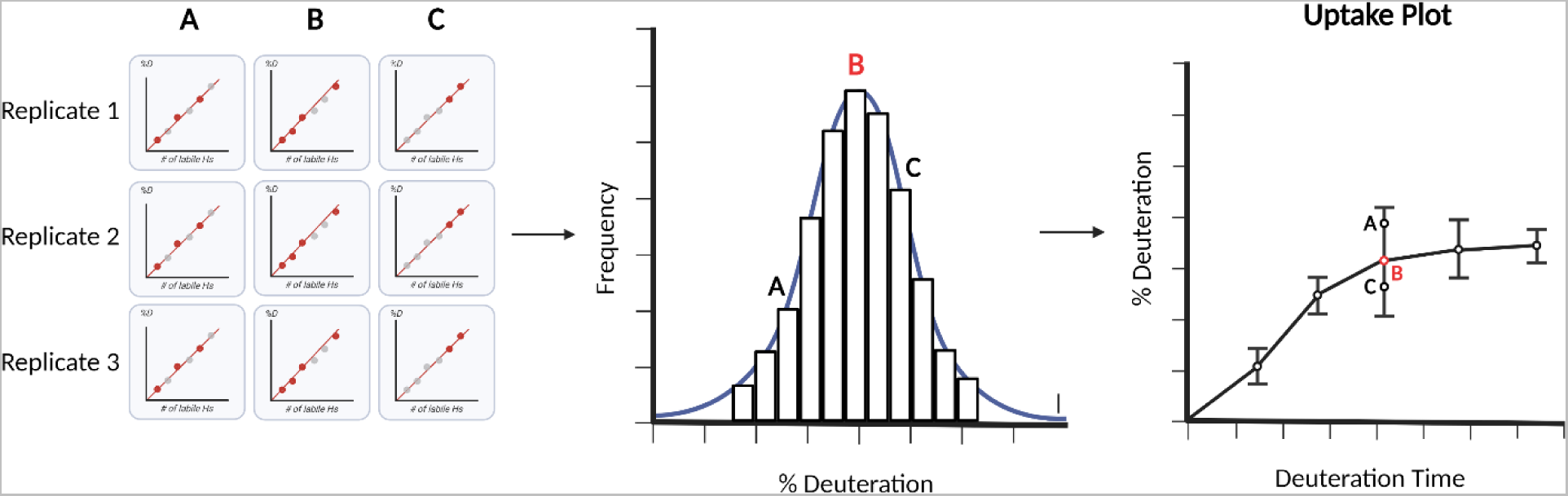
Data combination concept for uptake curves. After the selection of valid fragments and peptide distributions, shared across replicates and states, deuteration values calculated from all possible data combinations (illustrated as **A**, **B** and **C**) are used to generate a deuteration value distribution. The mean of the distribution is selected as the reported deuteration value, and the width of the distribution (95% CI) is used as an error bar on the uptake plot.

Mining high-redundancy data in this manner conveys two benefits. First, the fragment data validates the identity of the peptide because we require a minimum number of unique fragments. Second, the distribution tests the authenticity of the deuterium calculation. A peptide that produces a narrow distribution indicates an accurate and precise measure of deuteration, whereas a peptide that generates a non-normal and/or wide distribution highlights a compromised measurement. The flawed feature is then discarded from the dataset. Peptide deuteration kinetics from the phosphorylase B experiment were generated using this automation strategy and compared to a carefully curated manual analysis of the MS^1^ data (**Fig. 6**). The resulting heatmaps are almost indistinguishable, confirming that DIA-generated fragments can be used very effectively for automated peptide validation. The corresponding kinetics curves for all 380 peptides are provided in supplementary data (**Fig. S2**). During development, we detected a very slight bias against peptides with comparatively poor fragmentation, such as short singly charged peptides. To rescue high quality peptides in this category, we adopted a strategy from clinical mass spectrometry^40^. Qualifier transitions were required to validate peptide identity but were not used for deuteration calculations. Rather, they endorsed MS^1^-based deuteration measurements provided the latter were of high quality (*i.e.*, high isotopic fidelity).

**Figure 6.**
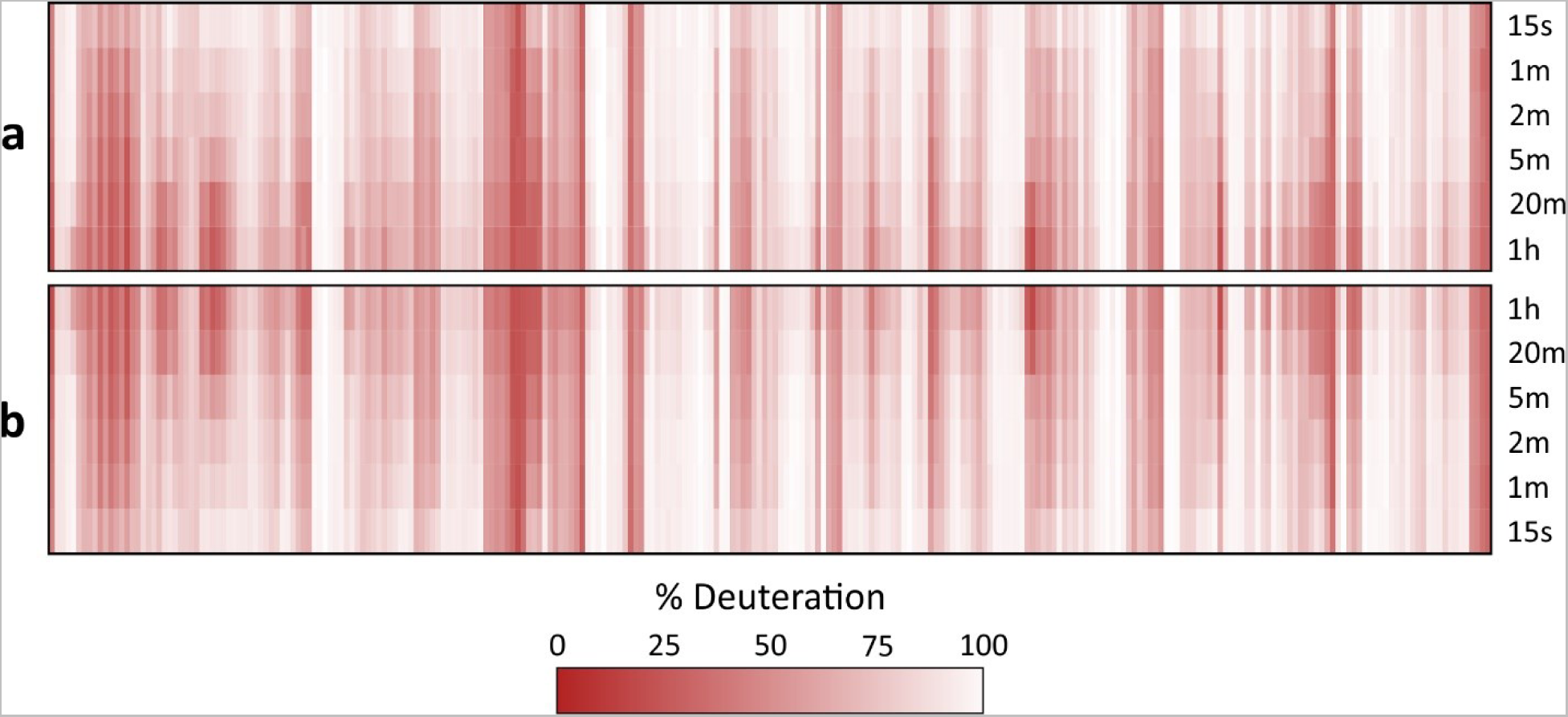
Comparison of manual MS1 and AutoHX-derived deuteration values. Data represents a 6-timepoint deuteration kinetics analysis of Phosphorylase B with all 380 peptide uptake plots shown as a heatmap ordered by sequence position. **(a) AutoHX** derived deuteration values as described in the text. (**b)** MS^1^-derived deuteration values. A minimum of three useful fragments per peptide shared across all replicates and timepoints was set as a requirement for a peptide to be accepted.

### Automated data authentication – differential analysis

HX-MS is most often used in a relational (or differential) manner. That is, deuteration kinetics for a protein in one state are compared to the same protein in a second state, either with a single labeling timepoint or an integration of the kinetic series. Common applications include drug or ligand characterization studies and quality control in the manufacture of protein biologics. These differential analyses are often depicted in a Woods plot, which shows induced changes in labeling as a function of protein sequence. To automate the generation of these plots, we developed a variation of the optimization method described above (**Fig. 5**). Here, the distribution is formed using the same combination strategy but the ΔD value is used instead. That is, given a specific combination of MS^1^ and MS^2^ data, average deuteration values are calculated from each replicate of a given state and compared to the average value from each replicate of the control state (**Fig. 7**). The distribution provides a solution to the problem of assigning significance to a given change in deuteration. The width of the distribution is used to assign a confidence interval to the change. An estimation of error overcomes the subjectivity of assigning relevance to a change based solely on its magnitude. This strategy provides a per-peptide measure of significance that has been lacking in the field, based on the subjectivity of manual data analysis. For example, a small change in a strongly ordered region of structure that takes up little deuterium can be classified as significant if the distribution is narrow. Peptides with conflicting deuteration values that cover areas of common sequence can be interpreted more rationally using the confidence interval as a guide.

**Figure 7:**
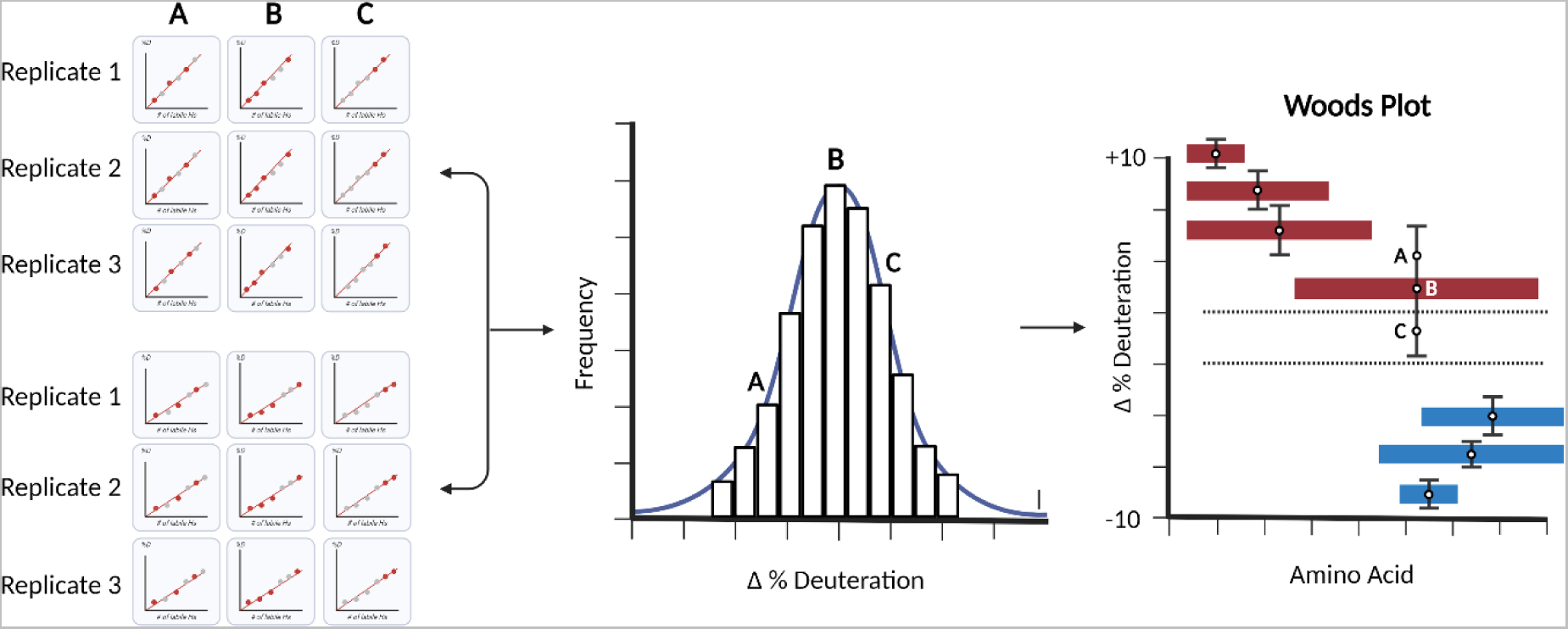
Data combination applied to differential analysis. Deuteration difference values between two states calculated from all possible data combinations (illustrated as **A**, **B** and **C**) are used to generate a deuteration difference value distribution. This establishes a sampling precision for any measured change and facilitates data interpretation. Red represents induced destabilizations and blue induced stabilizations.

### Differential analysis of Pol ϴ drug binding

To test this automation approach, we applied it to a manually validated dataset collected in DIA mode but analyzed in the conventional MS^1^-only fashion (manuscript accepted). Pol ϴ, a DNA polymerase, is a promising cancer drug target. It is upregulated in 70% of breast and epithelial ovarian cancers and it contributes to mechanisms of resistance to both conventional and emerging therapies^41^. The antibiotic novobiocin was recently shown to bind to the protein^42^. We generated a sequence map of Pol ϴ in the usual fashion and then conducted replicate HX-DIA analysis of novobiocin-bound vs free Pol ϴ. The deuterated data were analyzed in three ways. First, we naïvely applied the full sequence map and generated deuteration differences from the MS^1^ domain data. Second, the MS^1^ data were manually inspected by two experts and conflict-free peptides with good deuteration fit values were accepted. Third, AutoHX selected peptides and fragments automatically and determined the best subset to report. Woods plots were generated for each approach (**Fig. 8**).

**Figure 8.**
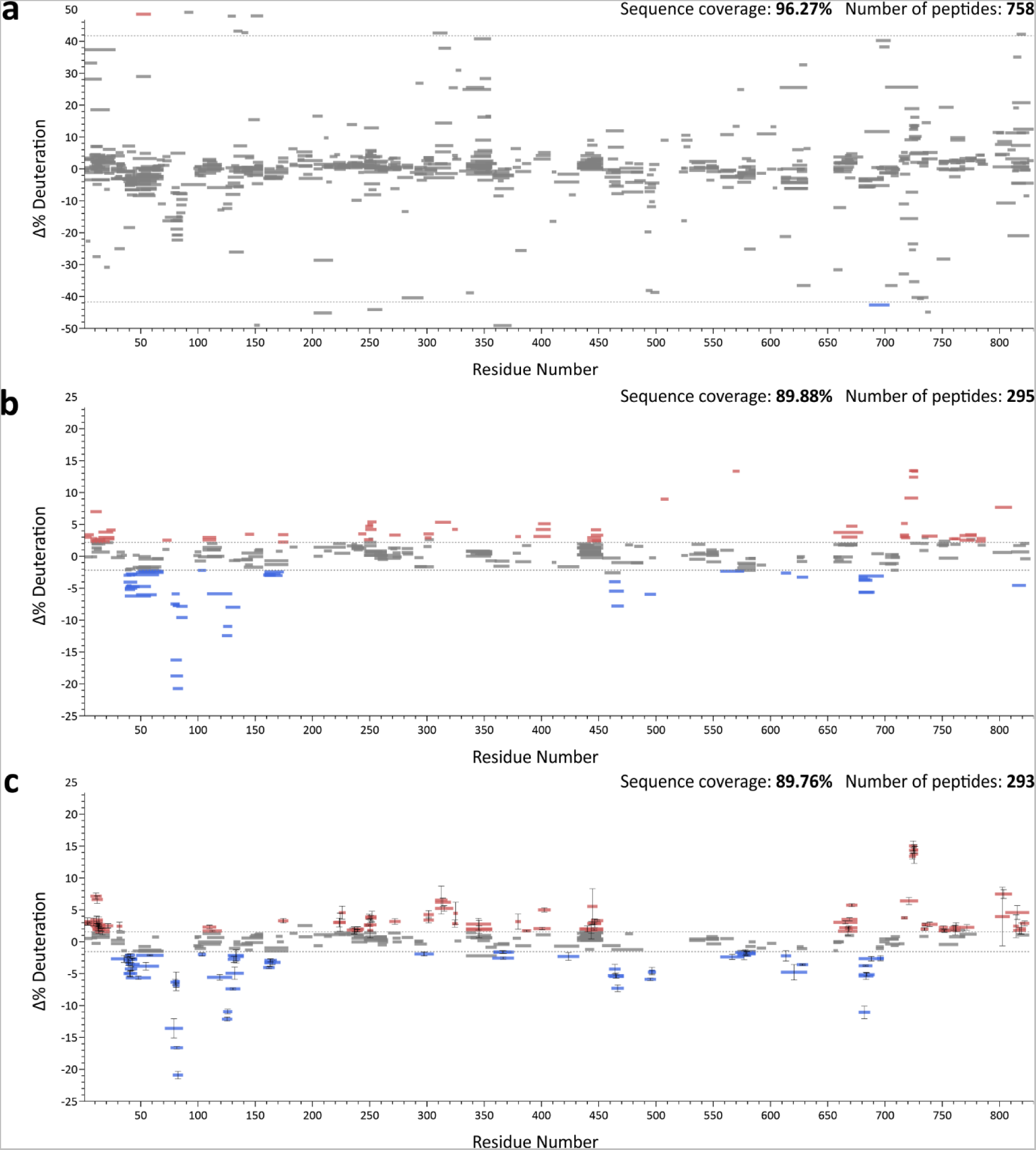
Directly comparing manual and automated analysis. (**a**) Unfiltered mapping file was blindly used for generation of a Woods plot for deuterated Pol ϴ data. (**b**) Corresponding Woods plot produced through expert manual validation of MS1 data (**c**) Automated generation of Woods plot in AutoHX using DIA data. Residue numbering is based on pdb:5AGA. Red represents induced destabilizations and blue induced stabilizations.

Not surprisingly, this exercise shows that validation is a key requirement in HX-MS analysis. Peptide IDs from a proteomics search do not guarantee good quality peptides in HX-MS analysis. Manual validation is required in all existing HX-MS software packages to remove outliers and other suspicious values caused by retention time misassignment and/or spectral overlap. Auto-validation using DIA data produces a map that is nearly identical to one generated from rigorous manual validation and indeed, it revealed some hidden biases in manual review (*e.g.*, assigning favorable changes in sequence regions already showing change). No manual input was needed apart from assigning initial values to processing parameters tied to data quality, such as ppm errors in MS^1^ and MS^2^ and retention time precision. We note that this specific Pol ϴ dataset was collected on a TOF instrument, which highlights the platform-independent nature of auto-validation routine. While not necessary or even encouraged, manual validation options are still retained in AutoHX.

### Differential analysis of drug binding to DNA-PKcs

Automating data analysis creates opportunities for applications that previously were highly impractical. For example, using affinity isolates as input for HX-MS is very appealing, as it would avoid recombinant protein production and difficulties in reconstituting functional states. Affinity pulldowns are typically low yielding and even with extensive washing, target proteins are often co-isolated with a significant fraction of nonspecific binding proteins.

To test the performance of DIA and AutoHX on such a challenging sample type, we analyzed DNA-PKcs in a microscale pulldown experiment and used the isolate as input to a drug binding analysis. DNA-PKcs is a protein kinase that regulates double-strand DNA break repair. It is also one of the largest mammalian proteins (~450kDa). It functions as a conformational switch at the point of commitment to the non-homologous end-joining repair pathway. Several experimental anticancer therapeutics target the ATP binding site, and only recently have their binding modes been modeled by cryo-EM^43^. The challenge, in part, involved isolation from over 100 L of cell culture equivalent to generate sufficient protein for analysis, given the difficulties associated with heterologous expression^43,44^. Here, we isolated GFP-tagged DNA-PKcs from the lysate from only two 10 cm plates of CHO cells, sufficient to generate enough material for the sequence map and six HX-MS experiments: three replicates of a drug-bound kinase and three ligand-free controls. We used AZD7648 as the drug, a selective inhibitor of DNA-PKcs that sensitizes cancer cells to radiation, doxorubicin and olaparib^45^.

We followed a typical HX-MS workflow with one exception. The higher complexity of the sample required a compositional analysis to build a searchable database and avoid false positive identifications. Using label free quantitation, we detected 30 proteins that represent at least 95% of the sample. DNA-PKcs itself contributed 30-35% of the total signal and produced 2414 peptides. We then naïvely generated a Woods plot from the six HX-MS experiments using the entire peptide list (**Fig. 9a**), revealing a complexity that would take days or weeks of manual validation to correct. AutoHX was able to process all six samples in 10 minutes on a single high-end desktop computer.

**Figure 9.**
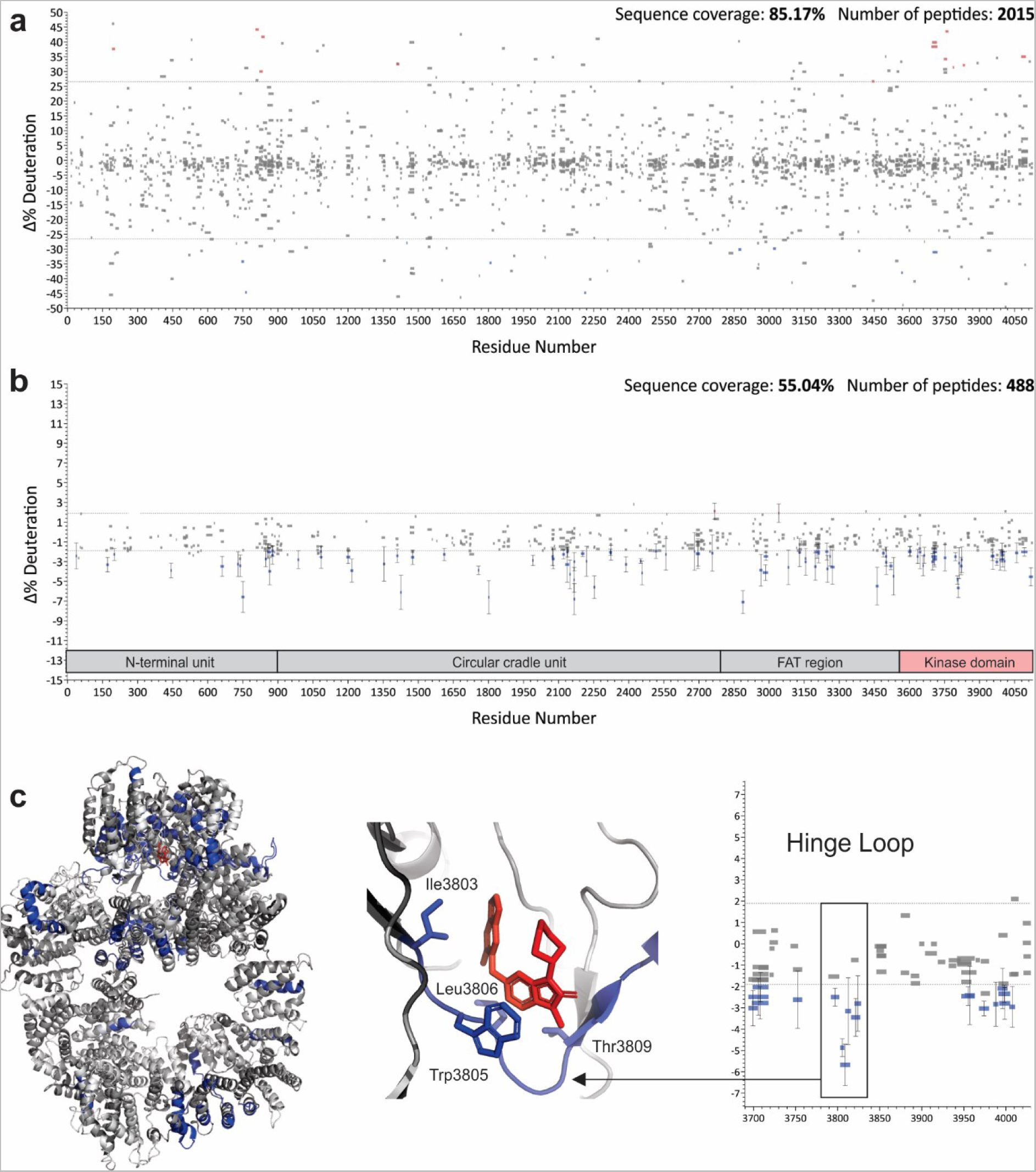
Accelerated HX-MS analysis of DNA-PKcs isolated from low quantities of CHO cells. (**a**) Unfiltered mapping file was blindly used for the generation of a Woods plot from replicate AZD7648-bound DNA-PKcs compared to a ligand-free control state. (**b**) Corresponding Woods plot produced using AutoHX without manual validation. (**c**) Mapping of high confidence stabilizations to PBB 7OTW, with expansion showing the highlighted hinge loop that defines the binding site. Key residues labeled as per Liang *et al.*^43^, where blue residues represent stabilization, dark grey no change and light grey no coverage.

The total number of usable peptides was reduced to 488, each with 4 or more unique fragments. This filter generated a sequence coverage of approximately 55% with strong coverage of the kinase domain, and widespread drug-induced stabilization of the protein is evident (**Fig. 9b**). This broad stabilization is anticipated. The control state is expected to be at least partially nucleotide-free and a previous HX analysis of the three-protein complex containing DNA-PKcs showed a similar effect arising from nucleotide binding^46^. The most confident changes were mapped onto the recent structure of AZD7648:DNA-PKcs, using only those peptides with error estimates outside of the noise limits (**Fig. 9c**). Interestingly, most of the detectable stabilizations are found in the FAT and kinase domains. One of the densest clusters identifies the hinge loop, which defines the primary binding site of the ligand^43^. Further optimization of the isolation should enhance this HX-MS assay and support the expansion of screening activities. We note that the conformational response of DNA-PKcs is critical to repair pathway commitment and is potentially druggable through allosteric inhibition. Accelerated HX-MS workflows should prove useful in exploring this concept.

## Conclusions

Automation tools have improved the rate at which HX-MS data can be collected, but the burden of manual data analysis has limited the extent to which the technology can be applied to many interesting problems. By invoking DIA methodology, AutoHX removes this burden while simultaneously providing both peptide validation and data authentication. There is a parallel to be drawn with the development history of proteomics. Early methods for protein detection relied on extensive fractionation (e.g. 2D gels) followed by fingerprint-based MS-only methods using MALDI TOF. The transition to MS/MS enabled the direct analysis of far more complex states, supported by complexity-tolerant search engines. HX-DIA provides a conceptually similar paradigm shift. It supports a truly proteomics-grade approach that should democratize a technology platform that has long been viewed as the domain of specialists.

## Acknowledgements

The authors would like to thank Dr. Kathy Meek, Michigan State University, for V3 cells stably expressing EGFP-DNA-PKcs. This work was funded by the Natural Sciences and Engineering Research Council of Canada Discovery Grants RGPIN 2017-04879 to DS and RGPIN-2019-04829 to SPLM. Figure design was done using BioRender.com for Figures 1, 3, 5 and 7 under Academic license.

