## Supporting Information for "Automating data analysis for hydrogen/deuterium exchange mass spectrometry using data-independent acquisition methodology"

**Contents:**

**Figure S1. RANSAC-based isotopologue filtering**

**Figure S2. Comparison of MS1-derived and AutoHX-derived deuteration values for individual peptides**

**Tutorial - AutoHX**

**AutoHX - Supplement:**

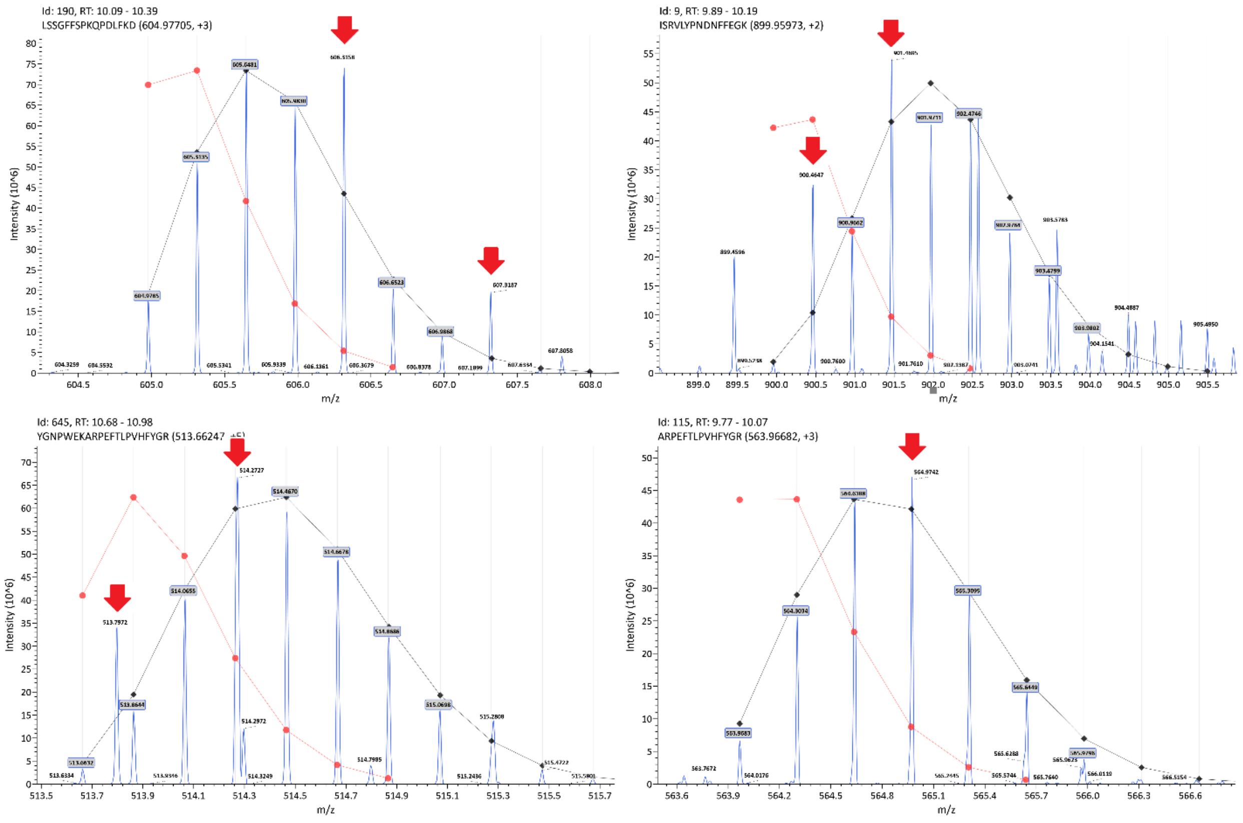


**Figure S1. RANSAC-based isotopologue filtering.** Peaks that deviate significantly from the deuteration model fit can be automatically unselected and not used for deuteration calculation. This is performed in both MS1 and MS2 data space to improve deuteration calculation. Examples are extracted from the Phosphorylase B kinetics dataset used in **Fig. 6** (main text).Red arrows indicate spurious peaks that the algorithm avoids selecting.

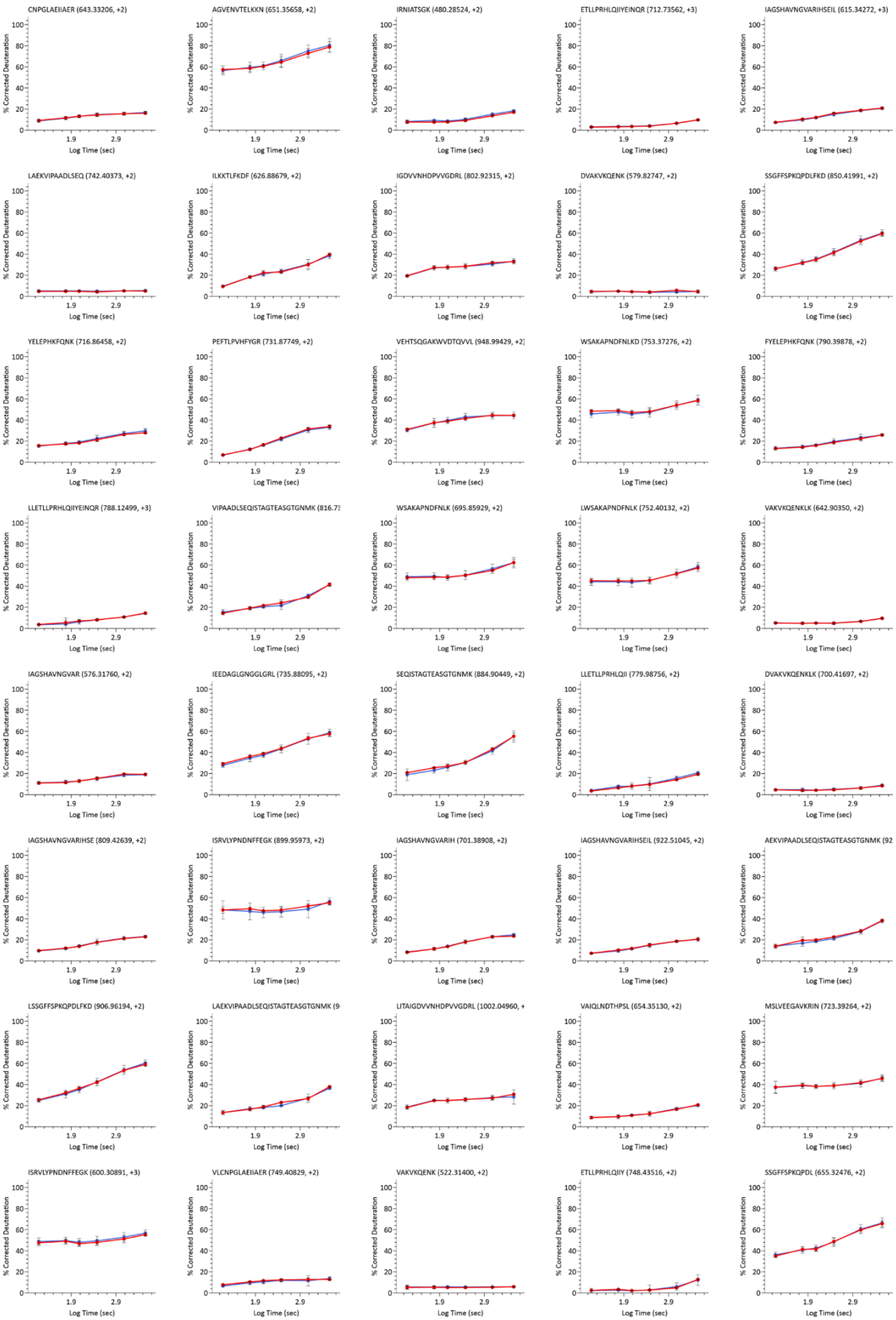
CNPGLAEIIAER (643.33206, +2)
AGVENVTELKKN (651.35658, +2)
IRNIATSGK (480.28524, +2)
ETLLPRHLQIIYEINQR (712.73562, +3)
IAGSHAVNGVARIHSEIL (615.34272, +3)
LAEKVIPAADLSEQ (742.40373, +2)
ILKKTLFKDF (626.88679, +2)
IGDVVNHDPVVGDRL (802.92315, +2)
DVAKVKQENK (579.82747, +2)
SSGFFSPKQPDLFKD (850.41991, +2)
YELEPHKFQNK (716.86458, +2)
PEFTLPVHFYGR (731.87749, +2)
VEHTSQGAKWVDTQVVL (948.99429, +2)
WSAKAPNDFNLKD (753.37276, +2)
FYELEPHKFQNK (790.39878, +2)
LLETLLPRHLQIIYEINQR (788.12499, +3)
VIPAADLSEQISTAGTEASGTGNMK (816.73
WSAKAPNDFNLK (695.85929, +2)
LWSAKAPNDFNLK (752.40132, +2)
VAKVKQENKLK (642.90350, +2)
IAGSHAVNGVAR (576.31760, +2)
IEEDAGLGNGGLGRL (735.88095, +2)
SEQISTAGTEASGTGNMK (884.90449, +2)
LLETLLPRHLQII (779.98756, +2)
DVAKVKQENKLK (700.41697, +2)
IAGSHAVNGVARIHSE (809.42639, +2)
ISRVLYPNDNFFEGK (899.95973, +2)
IAGSHAVNGVARIH (701.38908, +2)
IAGSHAVNGVARIHSEIL (922.51045, +2)
AEKVIPAADLSEQISTAGTEASGTGNMK (92
LSSGFFSPKQPDLFKD (906.96194, +2)
LAEKVIPAADLSEQISTAGTEASGTGNMK (9
LITAIGDVVNHDPVVGDRL (1002.04960, +
VAIQLNDTHPSL (654.35130, +2)
MSLVEEGAVKRIN (723.39264, +2)
ISRVLYPNDNFFEGK (600.30891, +3)
VLCNPGLAEIIAER (749.40829, +2)
VAKVKQENK (522.31400, +2)
ETLLPRHLQIIY (748.43516, +2)
SSGFFSPKQPDL (655.32476, +2)
% Corrected Deuteration
Log Time (sec)
0
20
40
60
80
100
1.9
2.9

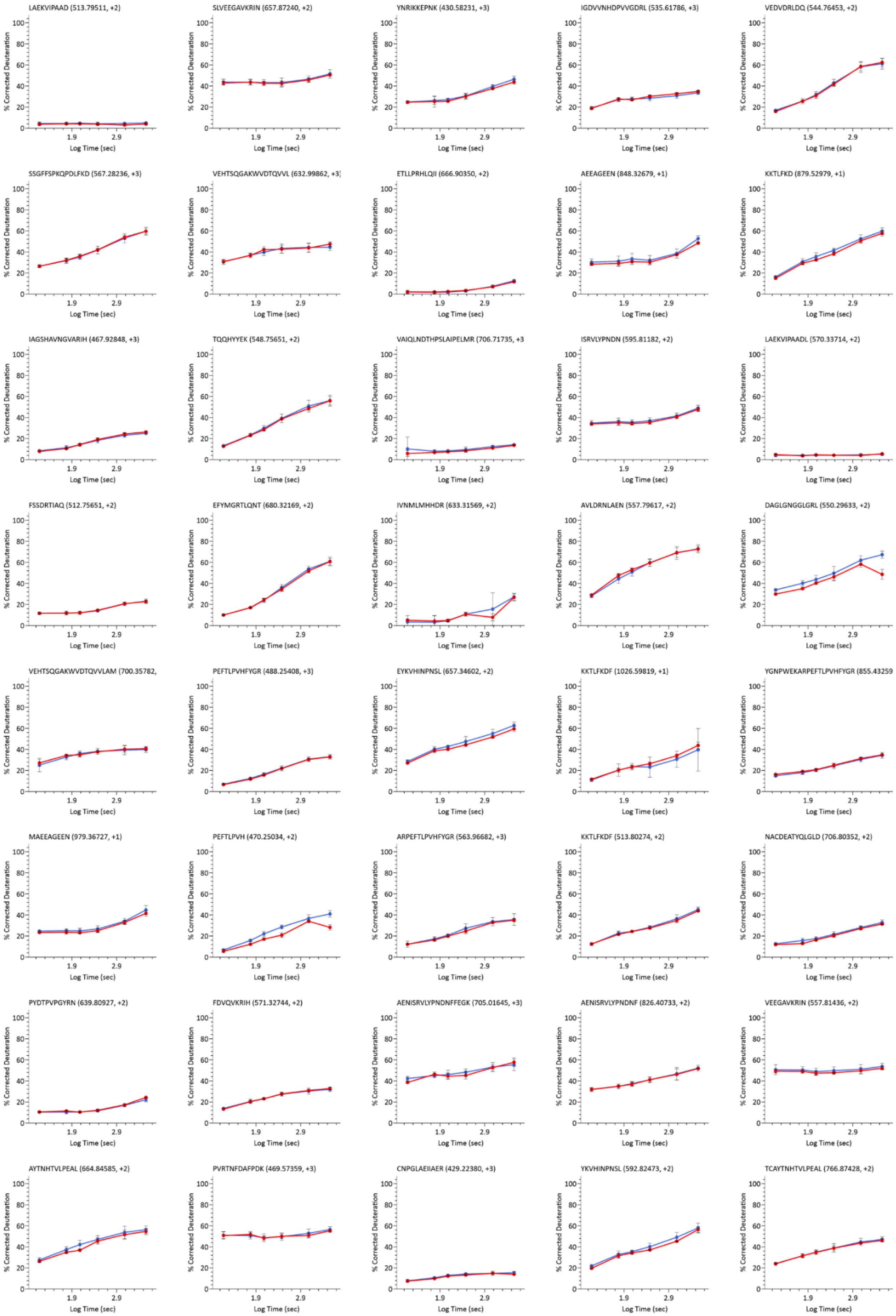
LAEKVIPAAD (513.79511, +2)
SLVEEGAVKRIN (657.87240, +2)
YNRIKKEPNK (430.58231, +3)
IGDVVNHDPVVGDRL (535.61786, +3)
VEDVDRLDQ (544.76453, +2)
SSGFFSPKQPDLFKD (567.28236, +3)
VEHTSQGAKWVDTQVVL (632.99862, +3)
ETLLPRHLQII (666.90350, +2)
AEEAGEEN (848.32679, +1)
KKTLFKD (879.52979, +1)
IAGSHAVNGVARIH (467.92848, +3)
TQQHYYEK (548.75651, +2)
VAIQLNDTHPSLAIPELMR (706.71735, +3
ISRVLYPNDN (595.81182, +2)
LAEKVIPAADL (570.33714, +2)
FSSDRTIAQ (512.75651, +2)
EFYMGRTLQNT (680.32169, +2)
IVNMLMHHDR (633.31569, +2)
AVLDRNLAEN (557.79617, +2)
DAGLGNGGLGRL (550.29633, +2)
VEHTSQGAKWVDTQVVLAM (700.35782,
PEFTLPVHFYGR (488.25408, +3)
EYKVHINPNSL (657.34602, +2)
KKTLFKDF (1026.59819, +1)
YGNPWEKARPEFTLPVHFYGR (855.43259
MAEEAGEEN (979.36727, +1)
PEFTLPVH (470.25034, +2)
ARPEFTLPVHFYGR (563.96682, +3)
KKTLFKDF (513.80274, +2)
NACDEATYQLGLD (706.80352, +2)
PYDTPVPGYRN (639.80927, +2)
FDVQVKRIH (571.32744, +2)
AENISRVLYPNDNFFEGK (705.01645, +3)
AENISRVLYPNDNF (826.40733, +2)
VEEGAVKRIN (557.81436, +2)
AYTNHTVLPEAL (664.84585, +2)
PVRTNFDAFPDK (469.57359, +3)
CNPGLAEIIAER (429.22380, +3)
YKVHINPNSL (592.82473, +2)
TCAYTNHTVLPEAL (766.87428, +2)
% Corrected Deuteration
Log Time (sec)

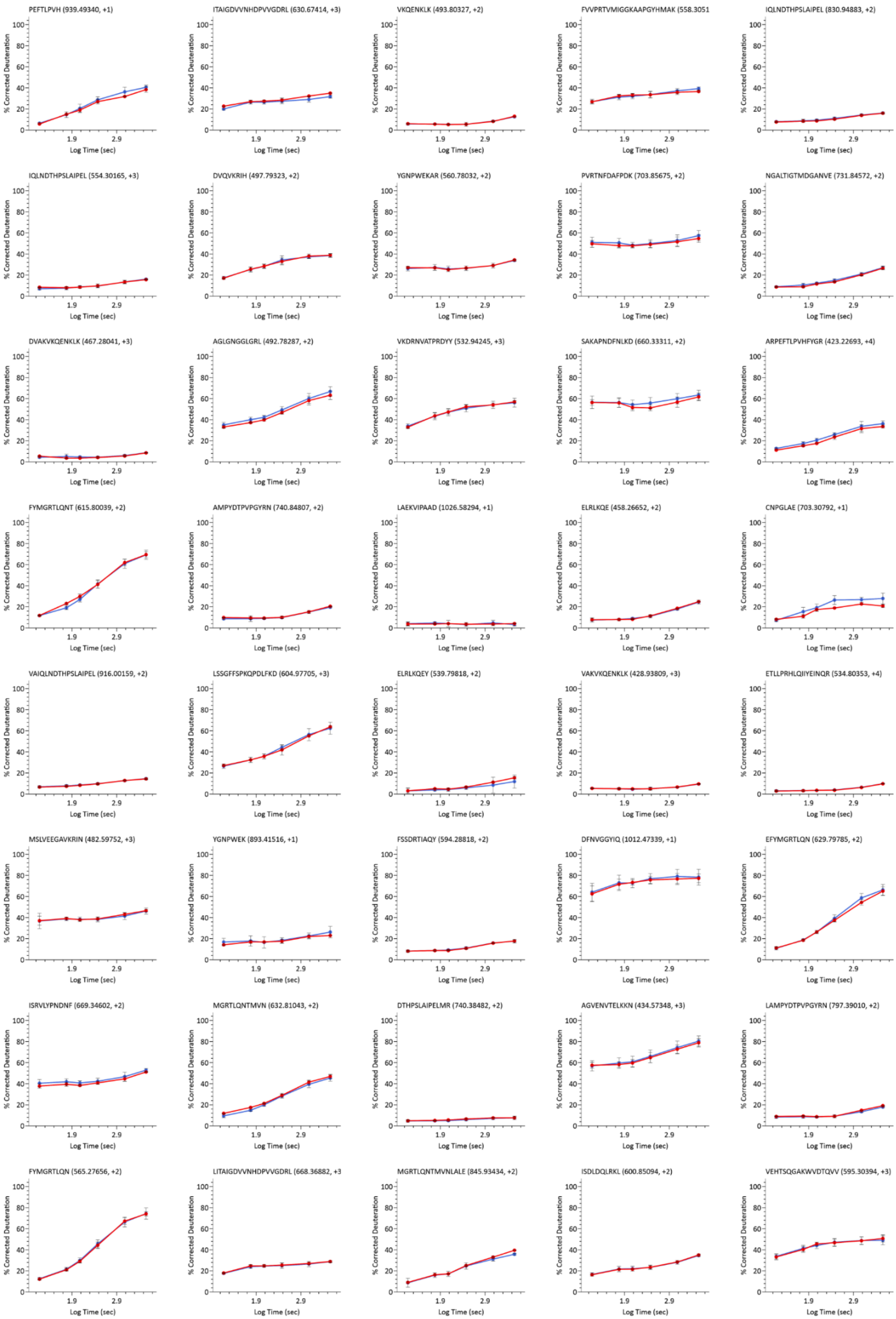
PEFTLPVH (939.49340, +1)
ITAIGDVVNHDPVVGDRL (630.67414, +3)
VKQENKLK (493.80327, +2)
FVVPRTVMIGGKAAPGYHMAK (558.3051
IQLNDTHPSLAIPEL (830.94883, +2)
IQLNDTHPSLAIPEL (554.30165, +3)
DVQVKRIH (497.79323, +2)
YGNPWEKAR (560.78032, +2)
PVRTNFDAFPDK (703.85675, +2)
NGALTIGTMDGANVE (731.84572, +2)
DVAKVKQENKLK (467.28041, +3)
AGLGNGGLGRL (492.78287, +2)
VKDRNVATPRDYY (532.94245, +3)
SAKAPNDFNLKD (660.33311, +2)
ARPEFTLPVHFYGR (423.22693, +4)
FYMGRTLQNT (615.80039, +2)
AMPYDTPVPGYRN (740.84807, +2)
LAEKVIPAAD (1026.58294, +1)
ELRLKQE (458.26652, +2)
CNPGLAE (703.30792, +1)
VAIQLNDTHPSLAIPEL (916.00159, +2)
LSSGFFSPKQPDLFKD (604.97705, +3)
ELRLKQEY (539.79818, +2)
VAKVKQENKLK (428.93809, +3)
ETLLPRHLQIIYEINQR (534.80353, +4)
MSLVEEGAVKRIN (482.59752, +3)
YGNPWEK (893.41516, +1)
FSSDRTIAQY (594.28818, +2)
DFNVGGYIQ (1012.47339, +1)
EFYMGRTLQN (629.79785, +2)
ISRVLYPNDNF (669.34602, +2)
MGRTLQNTMVN (632.81043, +2)
DTHPSLAIPELMR (740.38482, +2)
AGVENVTELKKN (434.57348, +3)
LAMPYDTPVPGYRN (797.39010, +2)
FYMGRTLQN (565.27656, +2)
LITAIGDVVNHDPVVGDRL (668.36882, +3
MGRTLQNTMVNLALE (845.93434, +2)
ISDLDQLRKL (600.85094, +2)
VEHTSQGAKWVDTQVV (595.30394, +3)
% Corrected Deuteration
Log Time (sec)

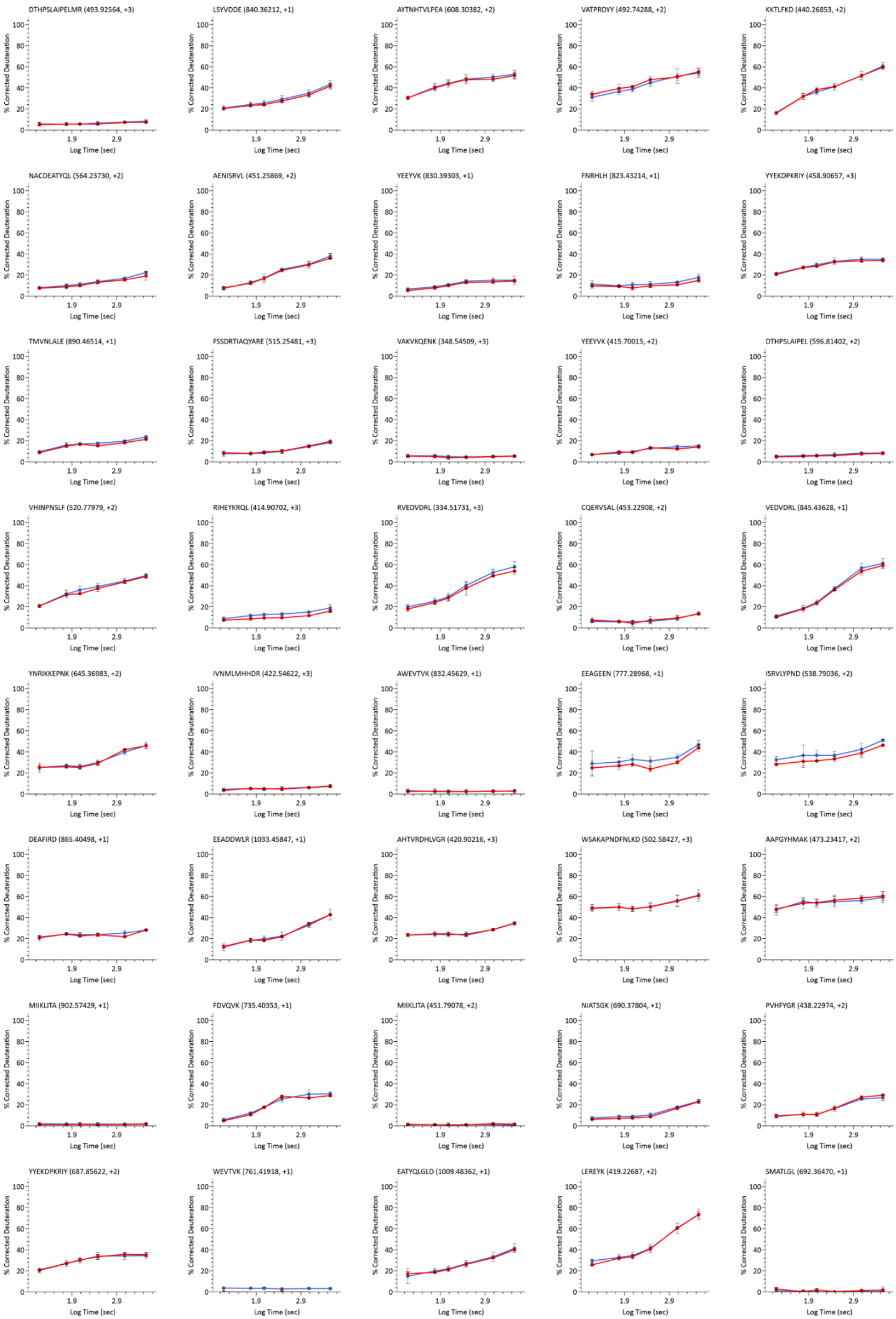
DTHPSLAIPELMR (493.92564, +3)
LSYVDDE (840.36212, +1)
AYTNHTVLPEA (608.30382, +2)
VATPRDYY (492.74288, +2)
KKTLFKD (440.26853, +2)
NACDEATYQL (564.23730, +2)
AENISRVL (451.25869, +2)
YEEYVK (830.39303, +1)
FNRHLH (823.43214, +1)
YYEKDPKRIY (458.90657, +3)
TMVNLALE (890.46514, +1)
FSSDRTIAQYARE (515.25481, +3)
VAKVKQENK (348.54509, +3)
YEEYVK (415.70015, +2)
DTHPSLAIPEL (596.81402, +2)
VHINPNSLF (520.77979, +2)
RIHEYKRQL (414.90702, +3)
RVEDVDRL (334.51731, +3)
CQERVSAL (453.22908, +2)
VEDVDRL (845.43628, +1)
YNRIKKEPNK (645.36983, +2)
IVNMLMHHDR (422.54622, +3)
AWEVTVK (832.45629, +1)
EEAGEEN (777.28968, +1)
ISRVLYPND (538.79036, +2)
DEAFIRD (865.40498, +1)
EEADDWLR (1033.45847, +1)
AHTVRDHLVGR (420.90216, +3)
WSAKAPNDFNLKD (502.58427, +3)
AAPGYHMAK (473.23417, +2)
MIIKLITA (902.57429, +1)
FDVQVK (735.40353, +1)
MIIKLITA (451.79078, +2)
NIATSGK (690.37804, +1)
PVHFYGR (438.22974, +2)
YYEKDPKRIY (687.85622, +2)
WEVTVK (761.41918, +1)
EATYQLGLD (1009.48362, +1)
LEREYK (419.22687, +2)
SMATLGL (692.36470, +1)
% Corrected Deuteration
Log Time (sec)

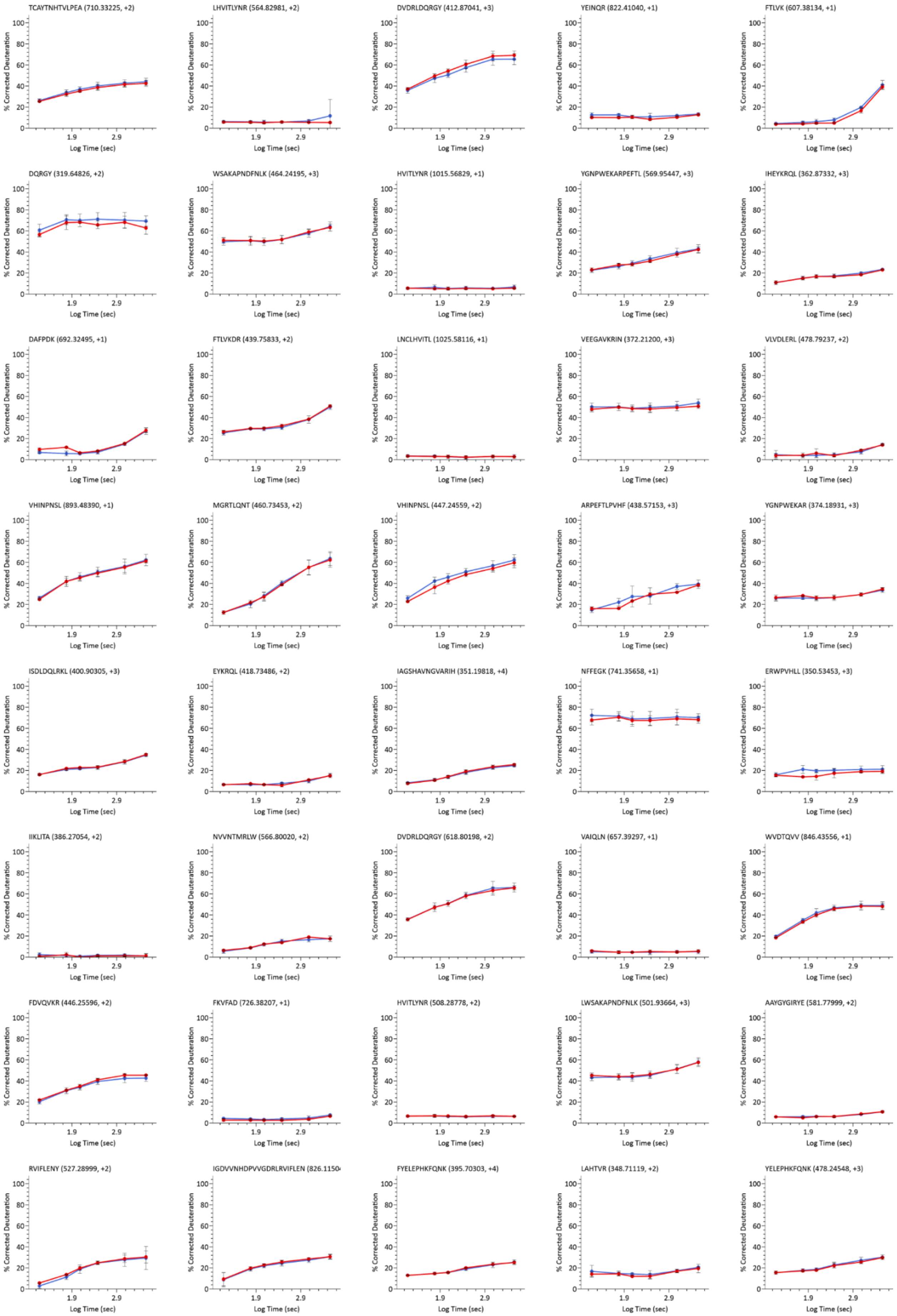
TCAYTNHTVLPEA (710.33225, +2)
LHVITLYNR (564.82981, +2)
DVDRLDQRGY (412.87041, +3)
YEINQR (822.41040, +1)
FTLVK (607.38134, +1)
DQRGY (319.64826, +2)
WSAKAPNDFNLK (464.24195, +3)
HVITLYNR (1015.56829, +1)
YGNPWEKARPEFTL (569.95447, +3)
IHEYKRQL (362.87332, +3)
DAFPDK (692.32495, +1)
FTLVKDR (439.75833, +2)
LNCLHVITL (1025.58116, +1)
VEEGAVKRIN (372.21200, +3)
VLVDLERL (478.79237, +2)
VHINPNSL (893.48390, +1)
MGRTLQNT (460.73453, +2)
VHINPNSL (447.24559, +2)
ARPEFTLPVHF (438.57153, +3)
YGNPWEKAR (374.18931, +3)
ISDLDQLRKL (400.90305, +3)
EYKRQL (418.73486, +2)
IAGSHAVNGVARIH (351.19818, +4)
NFFEGK (741.35658, +1)
ERWPVHLL (350.53453, +3)
IIKLITA (386.27054, +2)
NVVNTMRLW (566.80020, +2)
DVDRLDQRGY (618.80198, +2)
VAIQLN (657.39297, +1)
WVDTQVV (846.43556, +1)
FDVQVKR (446.25596, +2)
FKVFAD (726.38207, +1)
HVITLYNR (508.28778, +2)
LWSAKAPNDFNLK (501.93664, +3)
AAYGYGIRYE (581.77999, +2)
RVIFLENY (527.28999, +2)
IGDVVNHDPVVGDRLRVIFLEN (826.1150
FYELEPHKFQNK (395.70303, +4)
LAHTVR (348.71119, +2)
YELEPHKFQNK (478.24548, +3)
% Corrected Deuteration
Log Time (sec)

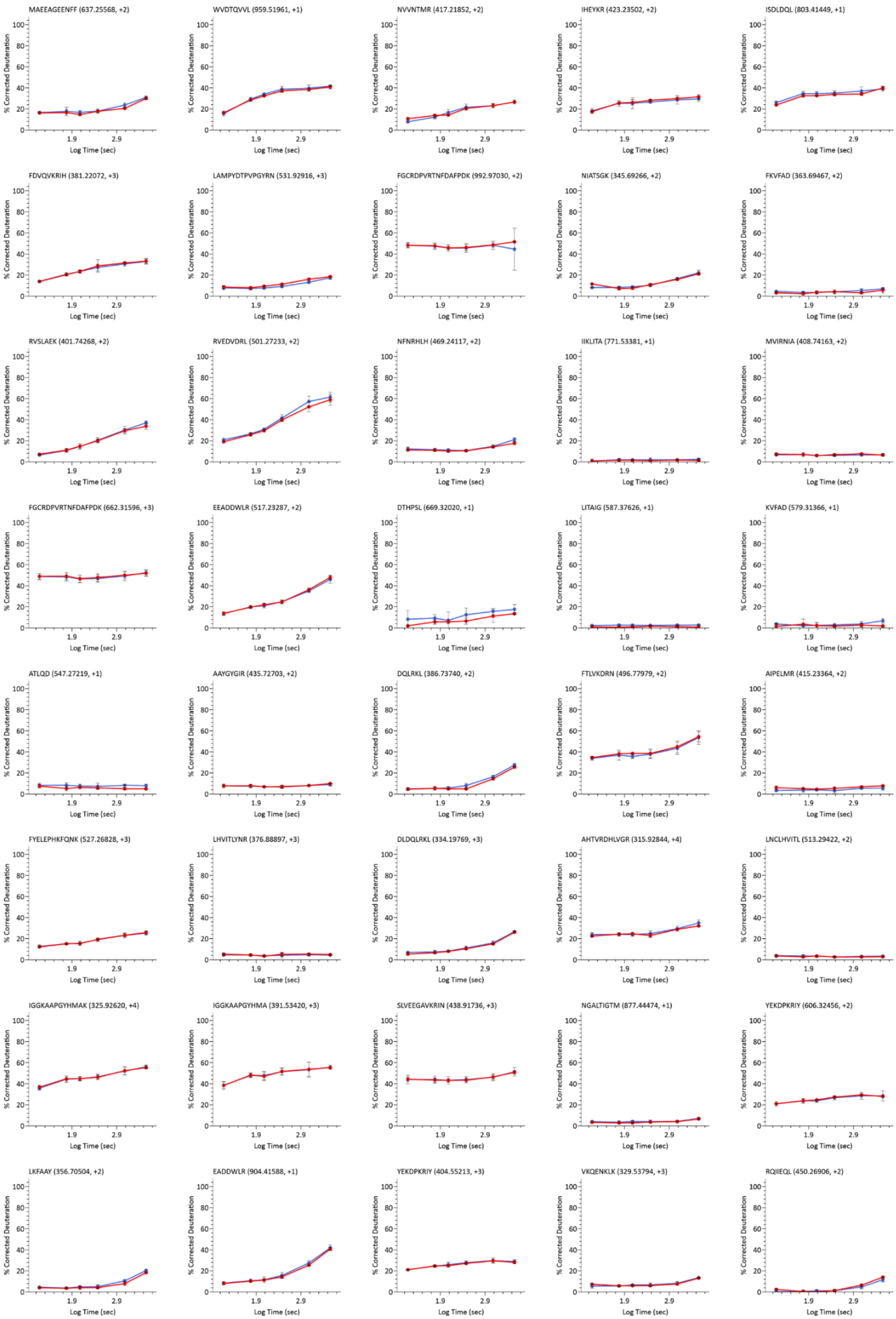
MAEEAGEENFF (637.25568, +2)
WVDTQVVL (959.51961, +1)
NVVNTMR (417.21852, +2)
IHEYKR (423.23502, +2)
ISDLDQL (803.41449, +1)
FDVQVKRIH (381.22072, +3)
LAMPYDTPVPGYRN (531.92916, +3)
FGCRDPVRTNFDAFPDK (992.97030, +2)
NIATSGK (345.69266, +2)
FKVFAD (363.69467, +2)
RVSLAEK (401.74268, +2)
RVEDVDRL (501.27233, +2)
NFNRHLH (469.24117, +2)
IIKLITA (771.53381, +1)
MVIRNIA (408.74163, +2)
FGCRDPVRTNFDAFPDK (662.31596, +3)
EEADDWLR (517.23287, +2)
DTHPSL (669.32020, +1)
LITAIG (587.37626, +1)
KVFAD (579.31366, +1)
ATLQD (547.27219, +1)
AAYGYGIR (435.72703, +2)
DQLRKL (386.73740, +2)
FTLVKDRN (496.77979, +2)
AIPELMR (415.23364, +2)
FYELEPHKFQNK (527.26828, +3)
LHVITLYNR (376.88897, +3)
DLDQLRKL (334.19769, +3)
AHTVRDHLVGR (315.92844, +4)
LNCLHVITL (513.29422, +2)
IGGKAAPGYHMAK (325.92620, +4)
IGGKAAPGYHMA (391.53420, +3)
SLVEEGAVKRIN (438.91736, +3)
NGALTIGTM (877.44474, +1)
YEKDPKRIY (606.32456, +2)
LKFAAY (356.70504, +2)
EADDWLR (904.41588, +1)
YEKDPKRIY (404.55213, +3)
VKQENKLK (329.53794, +3)
RQIIEQL (450.26906, +2)
% Corrected Deuteration
Log Time (sec)

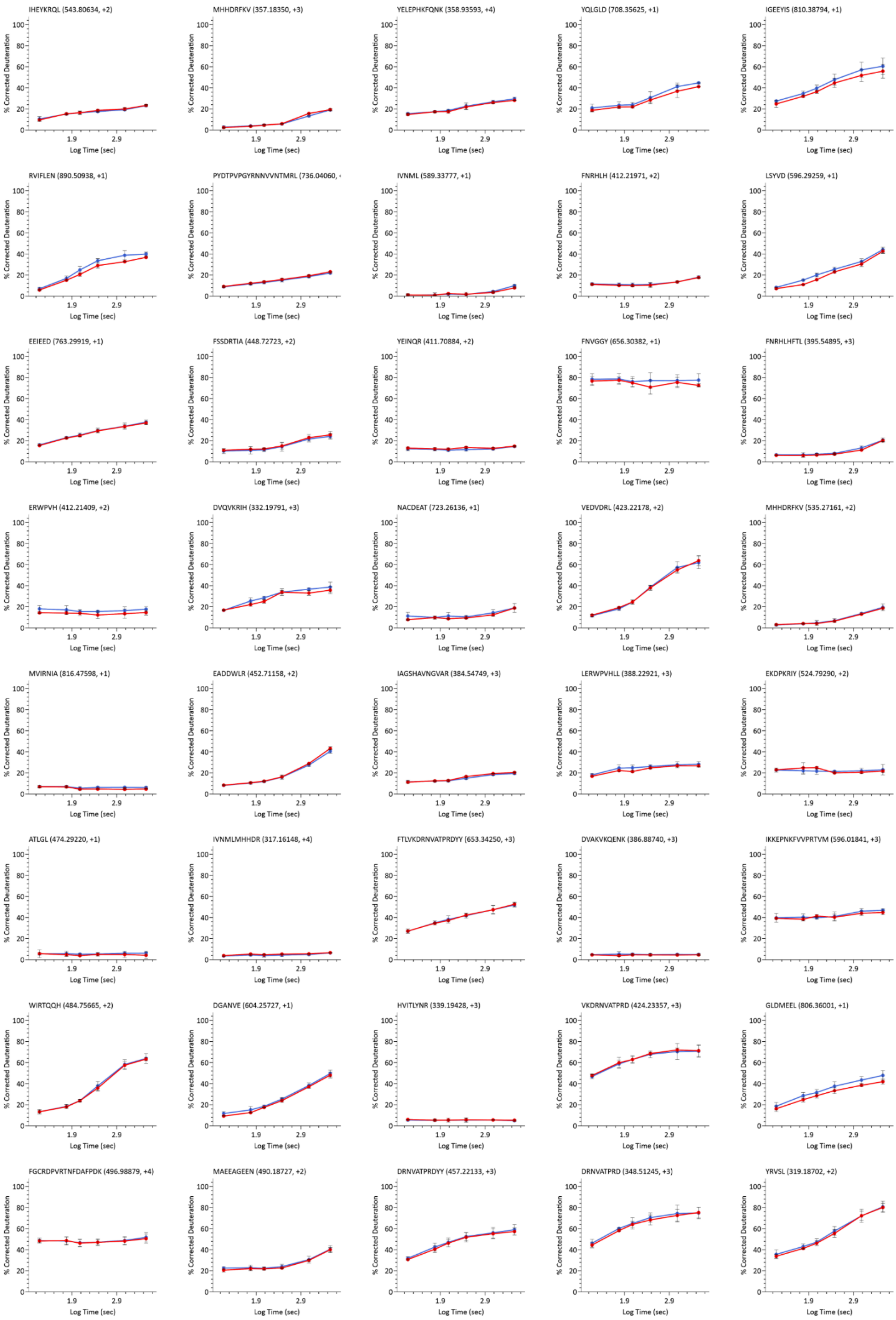

IHEYKRQL (543.80634, +2)
MHHDRFKV (357.18350, +3)
YELEPHKFQNK (358.93593, +4)
YQLGLD (708.35625, +1)
IGEEYIS (810.38794, +1)
RVIFLEN (890.50938, +1)
PYDTPVPGYRNNVVNTMRL (736.04060, +
IVNML (589.33777, +1)
FNRHLH (412.21971, +2)
LSYVD (596.29259, +1)
EEIEED (763.29919, +1)
FSSDRTIA (448.72723, +2)
YEINQR (411.70884, +2)
FNVGGY (656.30382, +1)
FNRHLHFTL (395.54895, +3)
ERWPVH (412.21409, +2)
DVQVKRIH (332.19791, +3)
NACDEAT (723.26136, +1)
VEDVDRL (423.22178, +2)
MHHDRFKV (535.27161, +2)
MVIRNIA (816.47598, +1)
EADDWLR (452.71158, +2)
IAGSHAVNGVAR (384.54749, +3)
LERWPVHLL (388.22921, +3)
EKDPKRIY (524.79290, +2)
ATLGL (474.29220, +1)
IVNMLMHHDR (317.16148, +4)
FTLVKDRNVATPRDYY (653.34250, +3)
DVAKVKQENK (386.88740, +3)
IKKEPNKFVVPRTVM (596.01841, +3)
WIRTQQH (484.75665, +2)
DGANVE (604.25727, +1)
HVITLYNR (339.19428, +3)
VKDRNVATPRD (424.23357, +3)
GLDMEEL (806.36001, +1)
FGCRDPVRTNFDAFPDK (496.98879, +4)
MAEEAGEEN (490.18727, +2)
DRNVATPRDYY (457.22133, +3)
DRNVATPRD (348.51245, +3)
YRVSL (319.18702, +2)
% Corrected Deuteration
Log Time (sec)

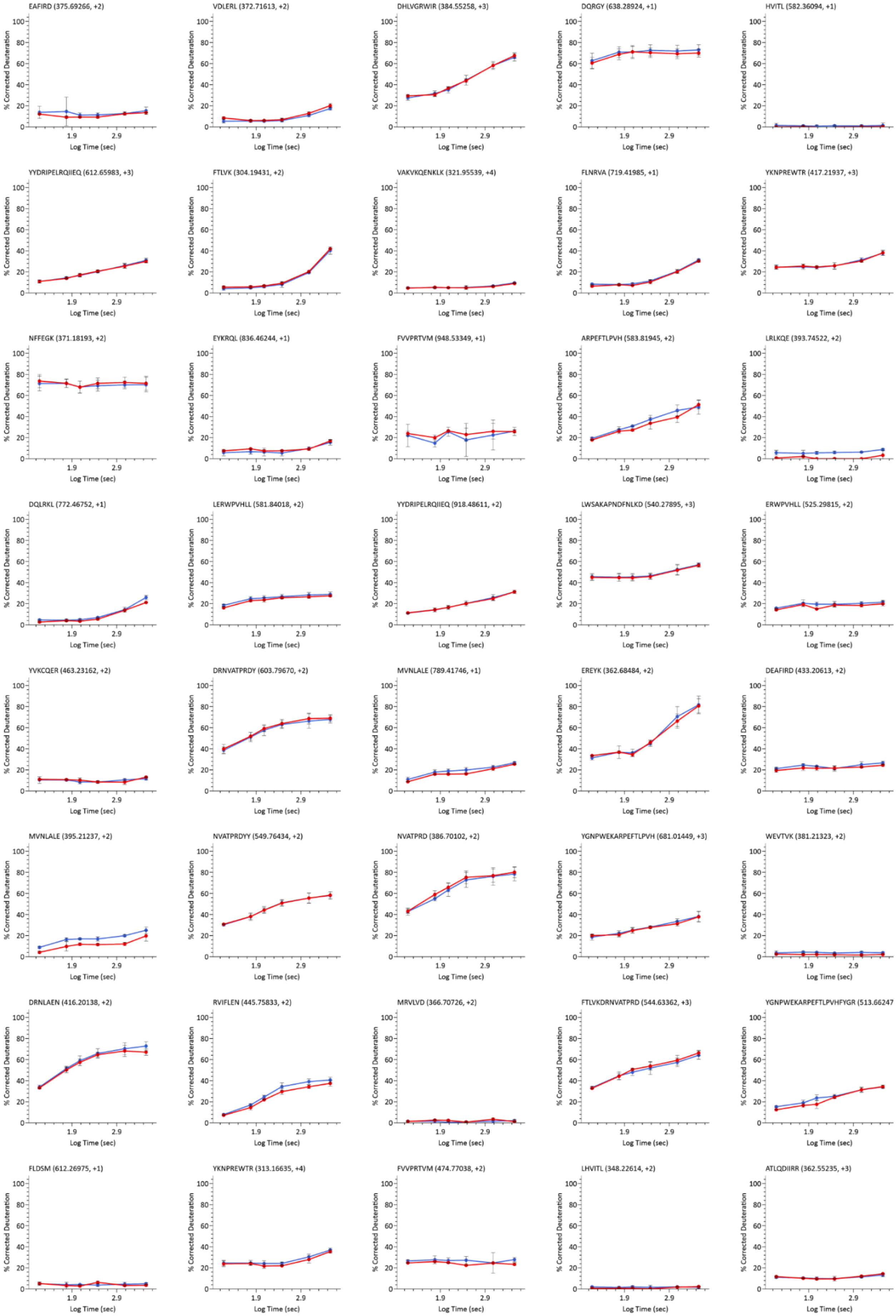
EAFIRD (375.69266, +2)
VDLERL (372.71613, +2)
DHLVGRWIR (384.55258, +3)
DQRGY (638.28924, +1)
HVITL (582.36094, +1)
YYDRIPELRQIIEQ (612.65983, +3)
FTLVK (304.19431, +2)
VAKVKQENKLK (321.95539, +4)
FLNRVA (719.41985, +1)
YKNPREWTR (417.21937, +3)
NFFEGK (371.18193, +2)
EYKRQL (836.46244, +1)
FVVPRTVM (948.53349, +1)
ARPEFTLPVH (583.81945, +2)
LRLKQE (393.74522, +2)
DQLRKL (772.46752, +1)
LERWPVHLL (581.84018, +2)
YYDRIPELRQIIEQ (918.48611, +2)
LWSAKAPNDFNLKD (540.27895, +3)
ERWPVHLL (525.29815, +2)
YVKCQER (463.23162, +2)
DRNVATPRDY (603.79670, +2)
MVNLALE (789.41746, +1)
EREYK (362.68484, +2)
DEAFIRD (433.20613, +2)
MVNLALE (395.21237, +2)
NVATPRDYY (549.76434, +2)
NVATPRD (386.70102, +2)
YGNPWEKARPEFTLPVH (681.01449, +3)
WEVTVK (381.21323, +2)
DRNLAEN (416.20138, +2)
RVIFLEN (445.75833, +2)
MRVLVD (366.70726, +2)
FTLVKDRNVATPRD (544.63362, +3)
YGNPWEKARPEFTLPVHFYGR (513.66247
FLDSM (612.26975, +1)
YKNPREWTR (313.16635, +4)
FVVPRTVM (474.77038, +2)
LHVITL (348.22614, +2)
ATLQDIIRR (362.55235, +3)
% Corrected Deuteration
Log Time (sec)

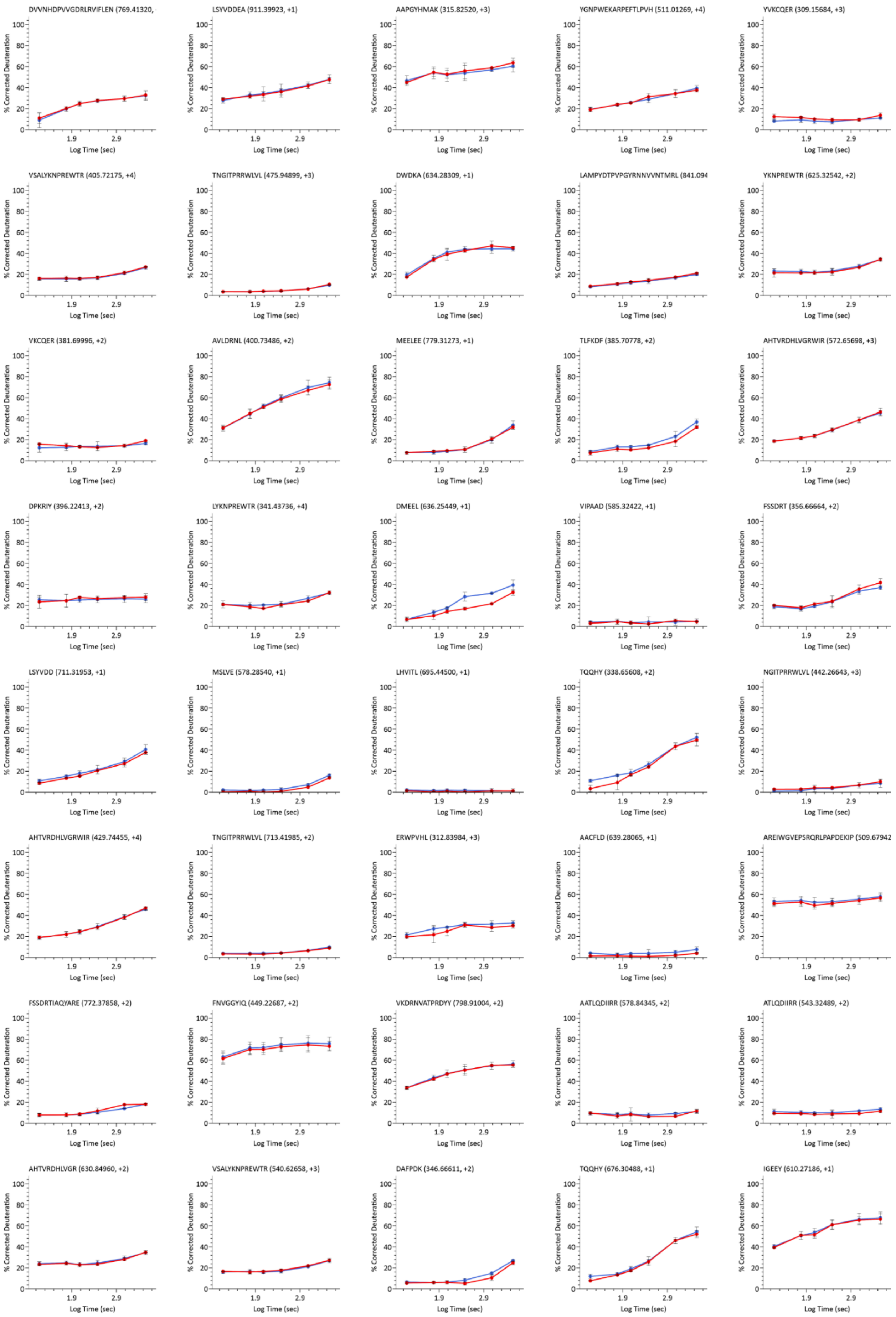
DVVNHDPVVGDRLRVIFLEN (769.41320,
LSYVDDEA (911.39923, +1)
AAPGYHMAK (315.82520, +3)
YGNPWEKARPEFTLPVH (511.01269, +4)
YVKCQER (309.15684, +3)
VSALYKNPREWTR (405.72175, +4)
TNGITPRRWLVL (475.94899, +3)
DWDKA (634.28309, +1)
LAMPYDTPVPGYRNNVVNTMRL (841.094
YKNPREWTR (625.32542, +2)
VKCQER (381.69996, +2)
AVLDRNL (400.73486, +2)
MEELEE (779.31273, +1)
TLFKDF (385.70778, +2)
AHTVRDHLVGRWIR (572.65698, +3)
DPKRIY (396.22413, +2)
LYKNPREWTR (341.43736, +4)
DMEEL (636.25449, +1)
VIPAAD (585.32422, +1)
FSSDRT (356.66664, +2)
LSYVDD (711.31953, +1)
MSLVE (578.28540, +1)
LHVITL (695.44500, +1)
TQQHY (338.65608, +2)
NGITPRRWLVL (442.26643, +3)
AHTVRDHLVGRWIR (429.74455, +4)
TNGITPRRWLVL (713.41985, +2)
ERWPVHL (312.83984, +3)
AACFLD (639.28065, +1)
AREIWGVEPSRQRLPAPDEKIP (509.67942
FSSDRTIAQYARE (772.37858, +2)
FNVGGYIQ (449.22687, +2)
VKDRNVATPRDYY (798.91004, +2)
AATLQDIIRR (578.84345, +2)
ATLQDIIRR (543.32489, +2)
AHTVRDHLVGR (630.84960, +2)
VSALYKNPREWTR (540.62658, +3)
DAFPDK (346.66611, +2)
TQQHY (676.30488, +1)
IGEEY (610.27186, +1)
% Corrected Deuteration
Log Time (sec)

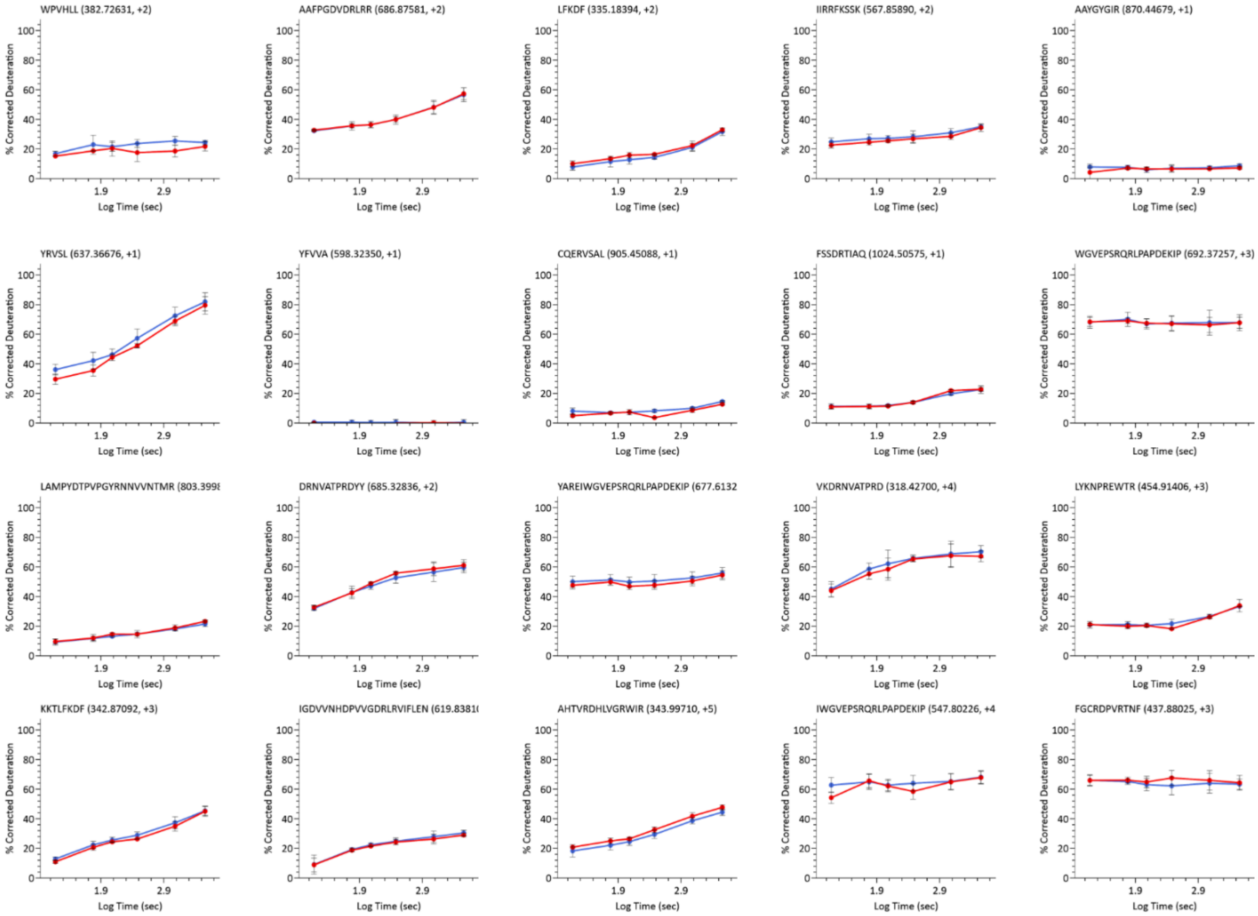


**Figure S2. Comparison of MS1-derived and AutoHX-derived deuteration values for individual peptides.** Kinetics plots from Phosphorylase B kinetics experiment showcasing manual $MS^1$-derived (Blue lines) and AutoHX-derived (Red lines) deuteration values. These values were used in the form of heatmap for **Fig. 6**, showing all 380 peptides.

### Tutorial -AutoHX for HX-DIA data analysis

A program is being established for early release and evaluation of the software. Interested individuals are encouraged to contact the corresponding author.  In anticipate of release, we make the following tutorial available:

#### Installation:

1. Install .NET Framework 3.5 SP1. For detailed instructions, visit this link.
2. [*Optional*] Install the latest version of Java 64-bit. This is required for some external tools (MS-GF+ and Percolator) used in HX-PIPE.
3. Run **setup.exe**. If you run the .msi directly, it will skip the check for additional system requirements. See below for more info.
4. During installation, you may be prompted to also install multiple "Visual C++ Redistributable 20XX" versions (2008, 2010, 2012, 2013, 2015+), if they are not already on your system. These pre-requisites are needed to make vendor software and other dependencies work. If you wish to install them manually, visit this link.

#### System requirements:

- Windows 7+ x64 with .NET 4.7.2 or higher.
- Multi-core CPU.
- 8GB+ RAM.
- 50GB+ of free disk space.

#### Create Project

Presently, AutoHX is a sub-module under the standard HX-DEAL workflow. Therefore, project creation for AutoHX in Mass Spec Studio follows the same overall wizard-based flow as regular HX-DEAL. The major difference is that DIA files are required to use the new HX-DIA processing routine and DIA bins need to be supplied for all the runs. Please follow these steps to get started:

1. **“New Project”:** Create a new HX-DEAL project by selecting “New Project” and selecting “HX-DEAL”. The default location for new projects will be inside your home directory. You can select a different location by clicking “Browse” and selecting a new directory.

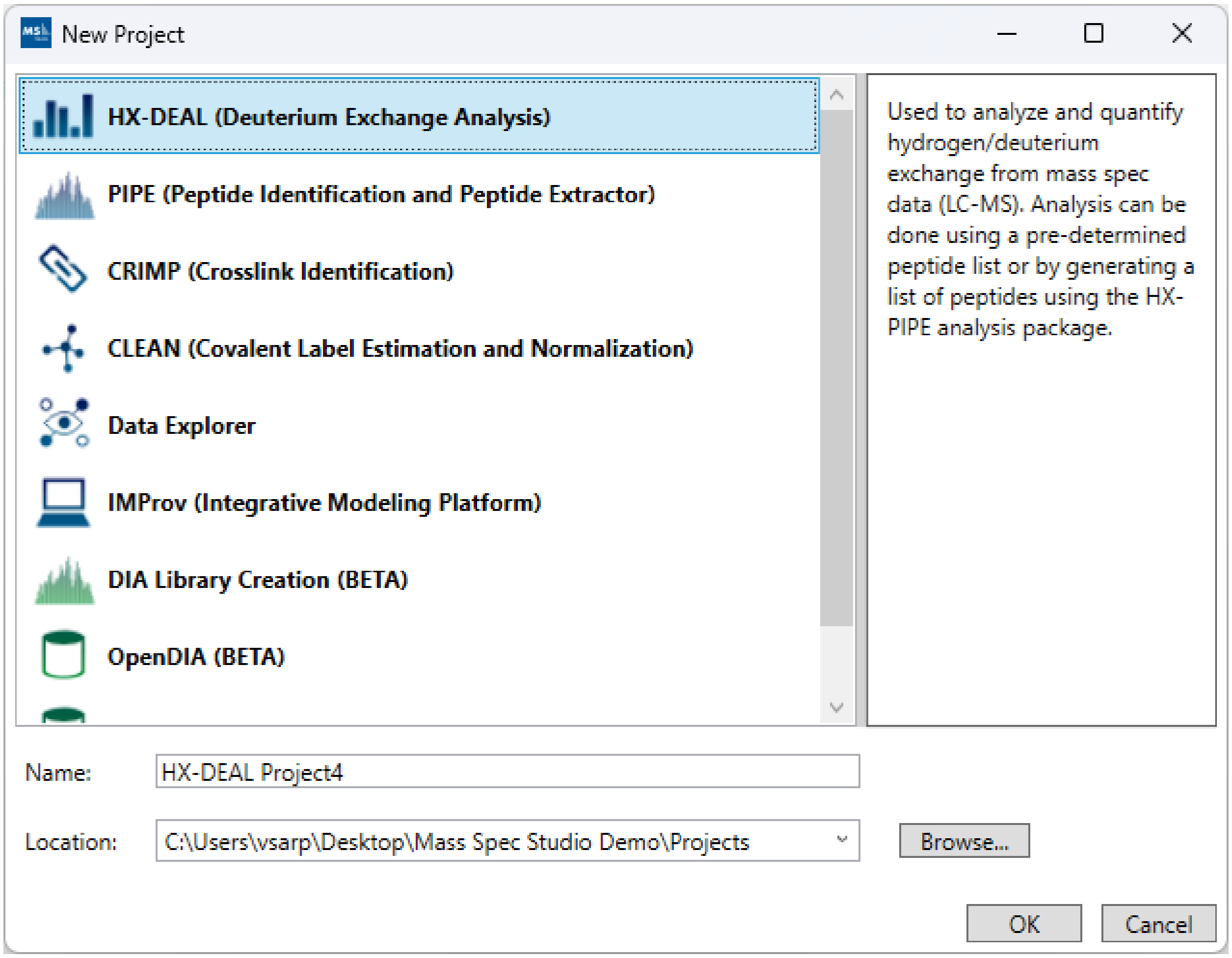


2. **"Add Proteins":** If you have more than a single protein or wish to use sequence visualizations, you can add your protein sequences either manually ("New"), via a FASTA file ("Select FASTA"), via a PDB file ("Select PDB(s)"), or via a 4-letter PDB code ("Fetch PDB(s)", example: "1jff"). Each protein will be listed as a separate row. If you have both PDB and FASTA, you can add the FASTA file first and then use the "Browse" button to link the PDB file. If you add both the PDB file and the FASTA file spearately, you may end up with duplicate proteins in the table. **Important:** For the sequence visualizations to work, the names of the proteins must match those found in the Peptide .csv file (explained later).
The "Merge polymers" option can be used to remove duplicate sequences from multimeric protein complexes which may appear as separate chains inside a PDB file. Example PDB-code: "1fu1".

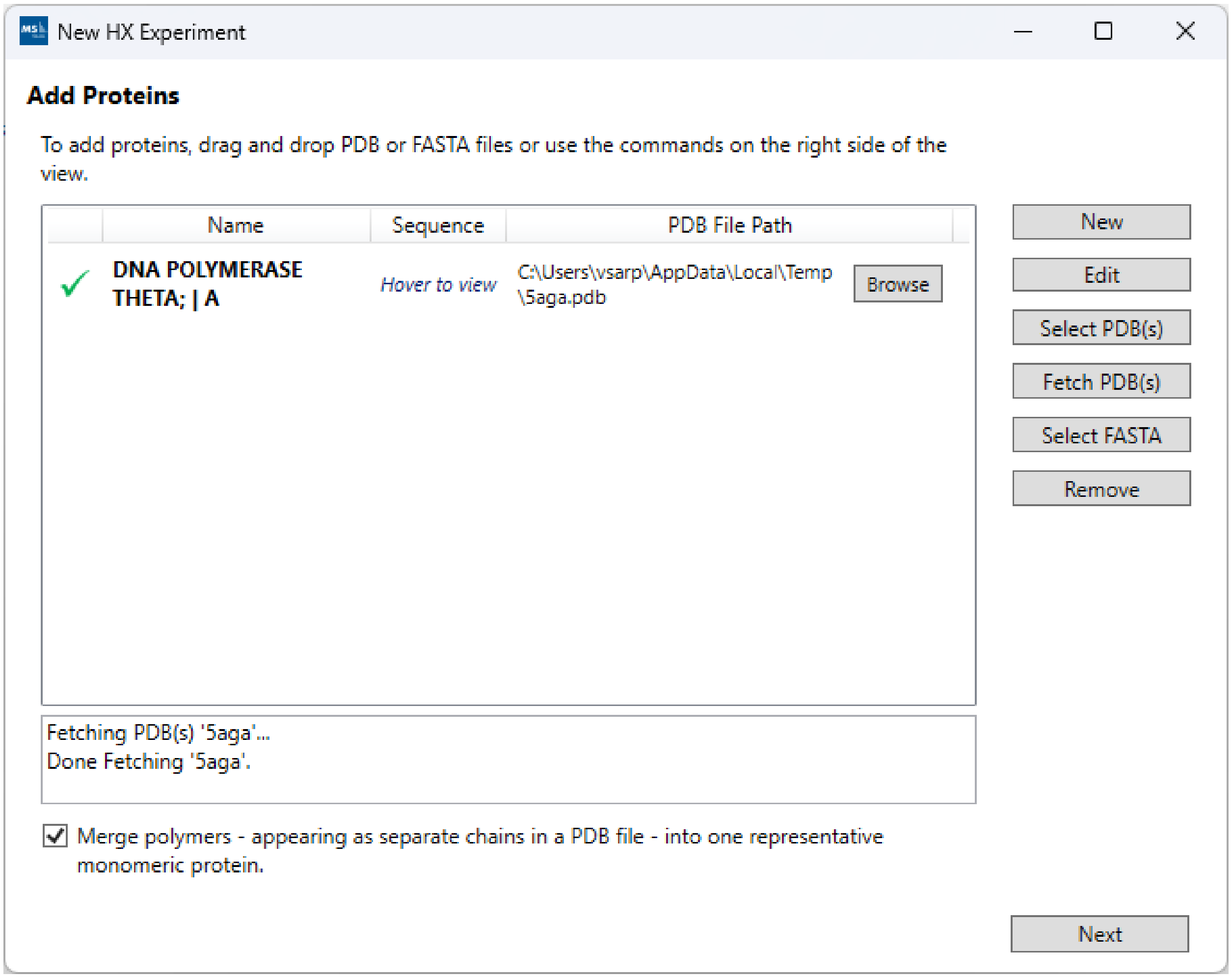


3. **“Add Protein States”:** For simple analysis, at least 1 protein state is required. For the additional comparative analysis tools and visualizations to become available, you must supply at least 2 proteins states.

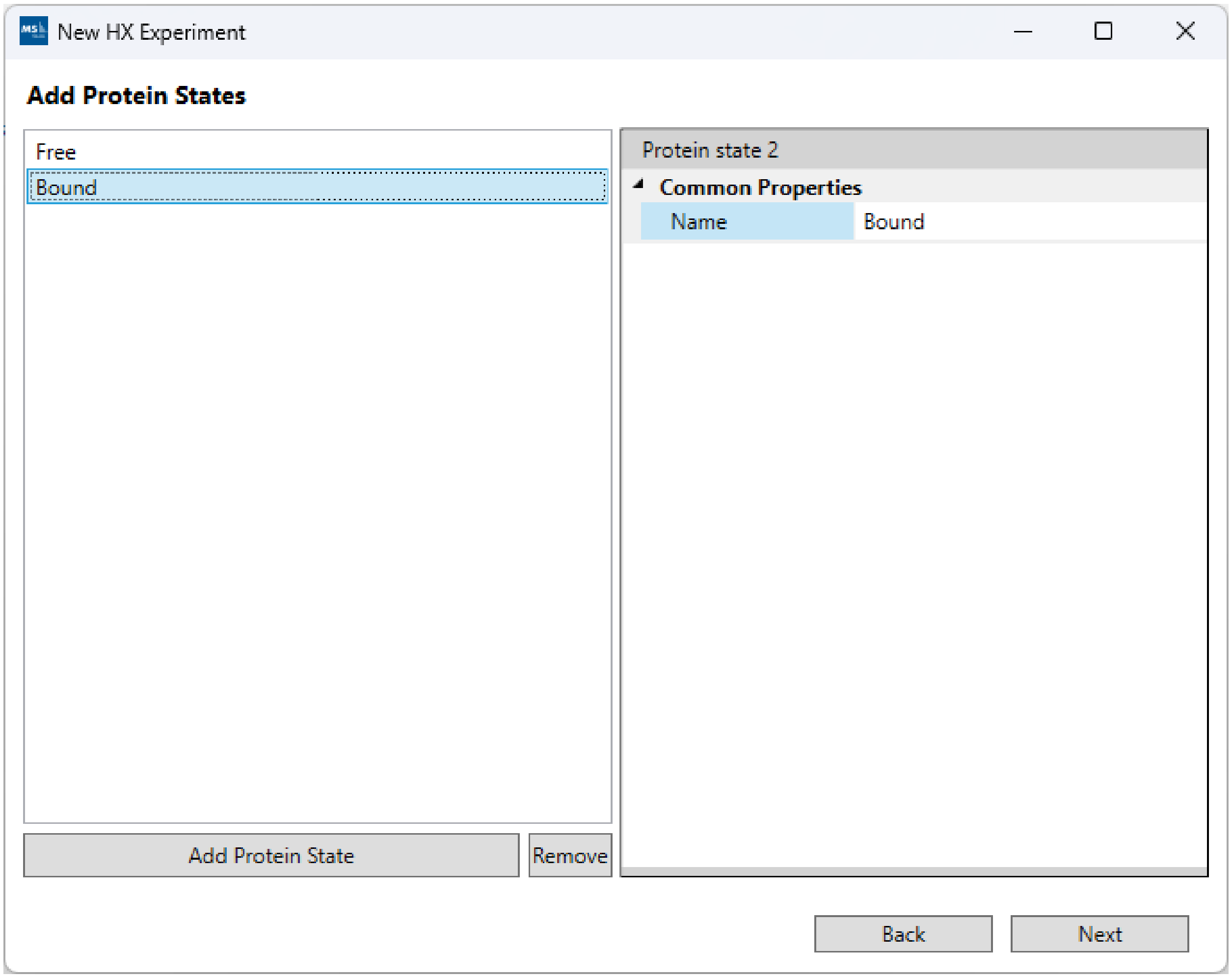


4. **“Add Labeling”:** At least 1 labeling condition is required (both time and %$D_2O$). For kinetics visualizations to become available, you must supply at least 2 labeling conditions.

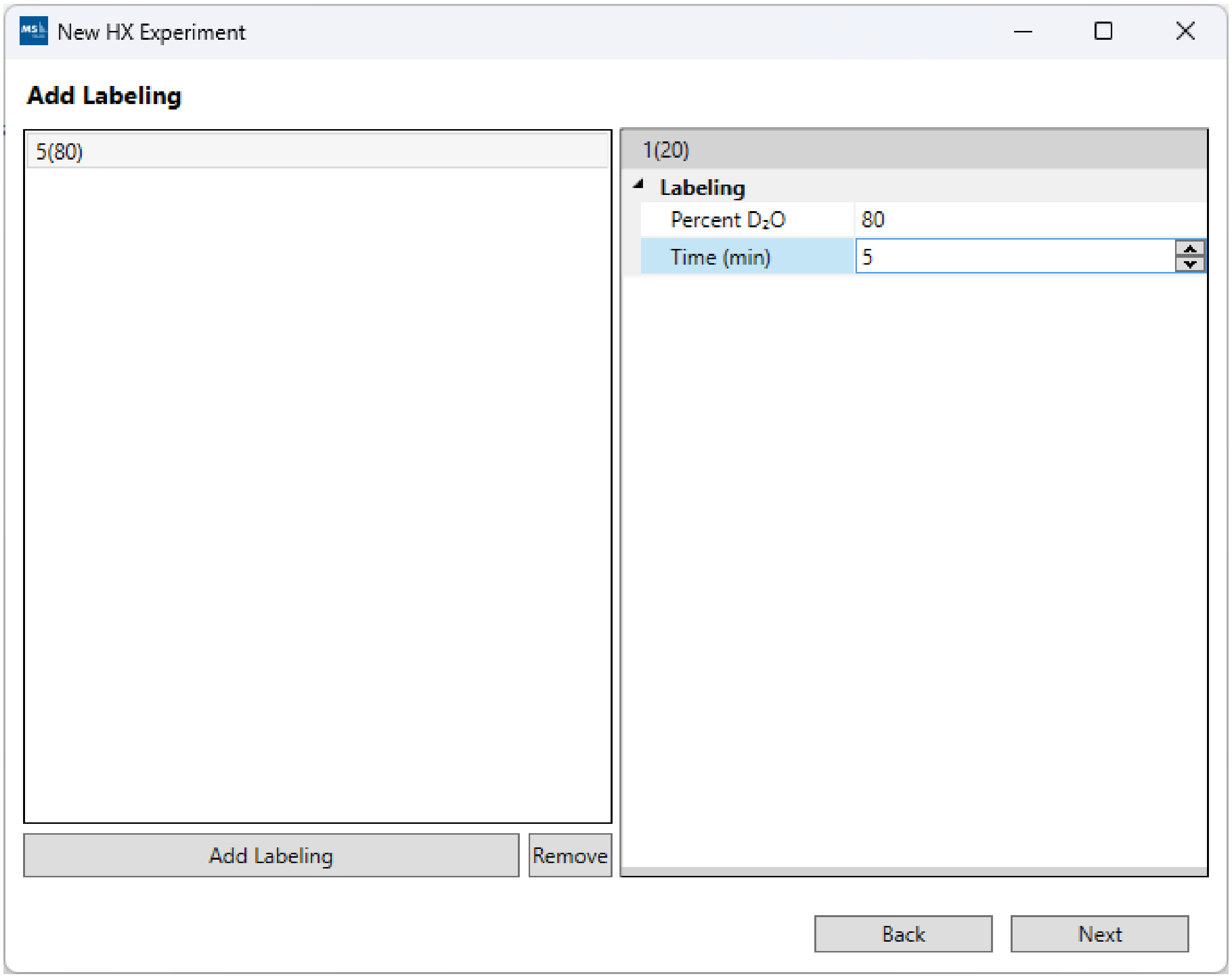


5. **"Add Runs":** Follow these steps to add your data:

a. Click "Browse" and select the root folder which contains your raw data files. In cases where the data files themselves are directories (example: Waters, Bruker), please make sure the containing folder is selected not the top-level .raw or .d directory itself. Note: Not all files have to be located under the same root folder from the start. After you add some files, you can still "Browse" to a different root directory and select additional files from the new location.

b. For most raw vendor files, <u>we strongly recommend to use "ProteoWizard Data Provider"</u>. If you already have Mass Spec Studio converted files (.mssdata, .mssmeta), select "Mass Spec Studio Data Provider".

c. We recommend you enable the "Convert to mssdata" checkbox. When ON, raw vendor files will be converted to the .mssdata format which enables super-fast searching at the cost of additional disk space for the .mssdata files. You can still proceed without converting the files, but the processing will be very slow. The "Noise filter" value is a multiplier of the minimum signal in each spectrum. A noise filter of “2” will remove any intensities smaller than 2 * mininum. A value of “0” will not remove any data.

d. Select your replicates (shift+click or ctrl+click for multi-select), select the appropriate Protein State/Labeling node and click the “>” button. If you make mistakes, you can remove runs using the “<” button.

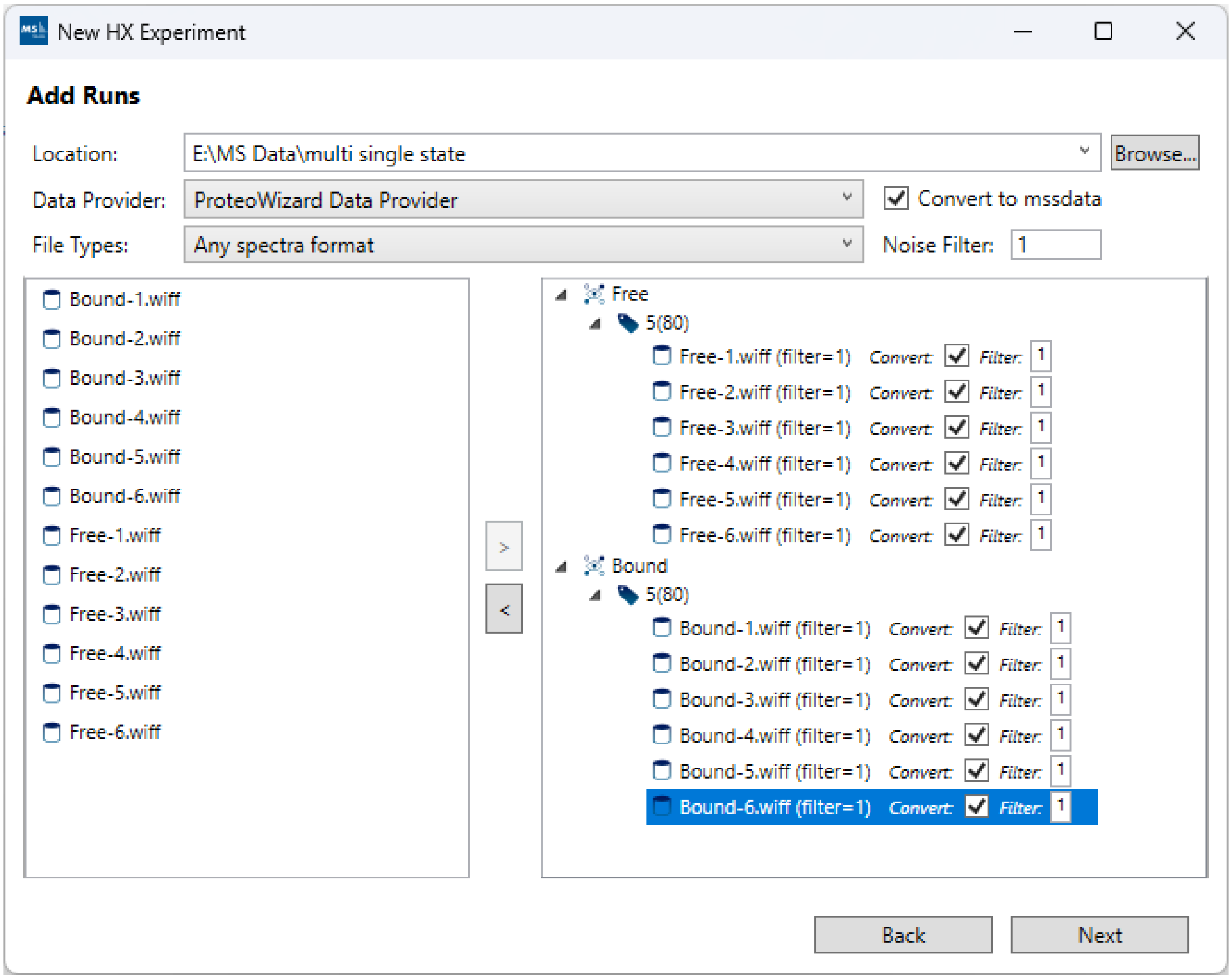


6. **“Configure Runs”:** For DIA files, select the “MSMS Data Independent Acquisition (DIA)” option. This will unlock a new column in the runs table for adding fragmentation bins (isolation window) for each run. The bins should be in “.csv” format with “Start” and “End” columns which represent the boundaries of each DIA isolation window. If all runs have the same isolation windows, the file can be supplied once under the first row representing “All” runs.

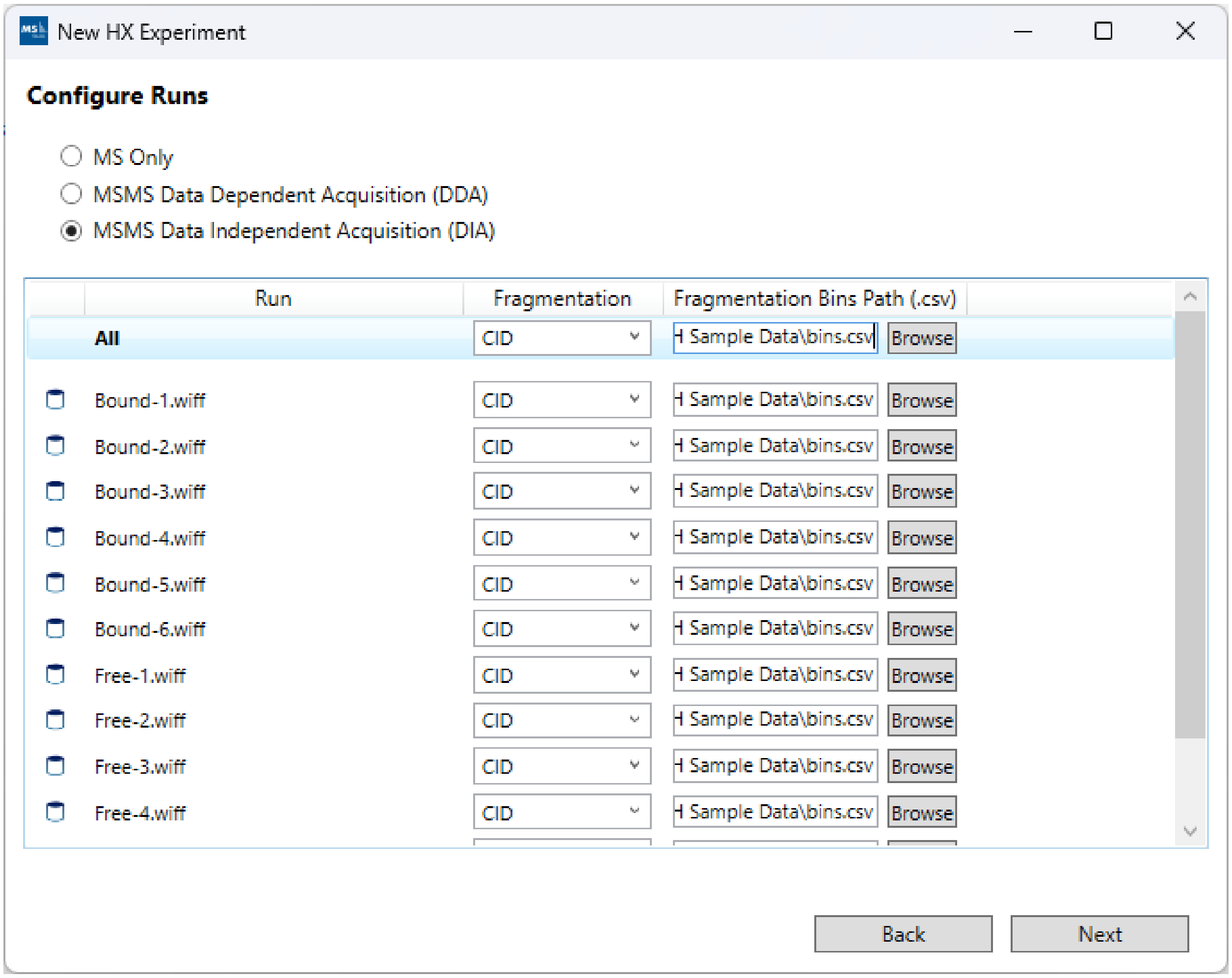


7. **"Provide a pre-identified peptide list (.csv)":** Click "Browse" and select a peptide list. Note: To avoid duplicates, it's recommended that your peptide identifications are grouped by sequence and charge such that each peptide (sequence/charge pair) has only one "RT". If you start with multiple separate peptide-spectrum matches (PSMs) for the same peptide, you can group them by sequence/charge and average their RTs to get the final peptide "RT". If you expect high discrepancies between the mapping runs and the DIA runs, you can define a larger "RT Variance" column in your peptides file. HX-DIA does not need a pre-selected transitions file because it will check all possible fragments for each peptide. However, if you wish to constrain the deuteration calculations to a pre-selected list of fragments, please supply them in the pre-selected transitions file.

A sample peptide list: https://s3.us-west-2.amazonaws.com/www.msstudio.ca/assets/tutorials/hx-deal/sample_peptides.csv

A sample transitions list (optional): https://s3.us-west-2.amazonaws.com/www.msstudio.ca/assets/tutorials/hx-deal/sample_transitions.tsv

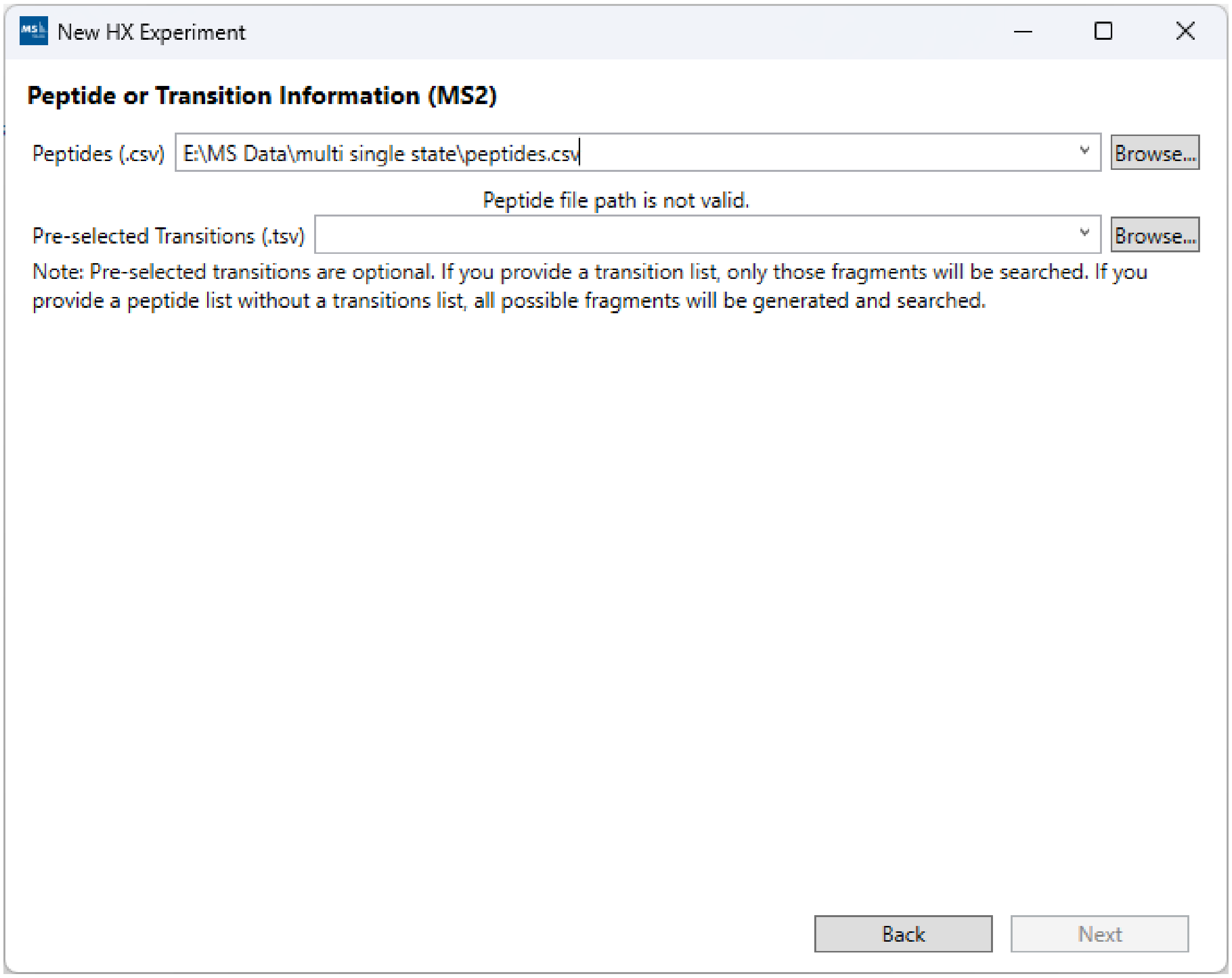


**Processing**

In order to use the HX-DIA processing routine (core of AutoHX), you must first enable "Beta Features" from the Tool -> Performance Manager window. A password is required to enable the AutoHX features – please contact us to obtain one.

To get started with processing, you can open the processing window from the Process menu and select the "Peptide HX-DIA" routine. The most important parameters to set correct are the mass accuracy and the peptide elution time parameters. The rest of the parameters are already set to the default values we found most useful during our internal testing. For advanced users, the advanced parameters to fine-tune the search can be enabled via the top-right "Advanced Parameters" checkbox. If you wish to know more about the parameters, you can click on the help tooltip buttons ("?") next to each processing step/subsection.

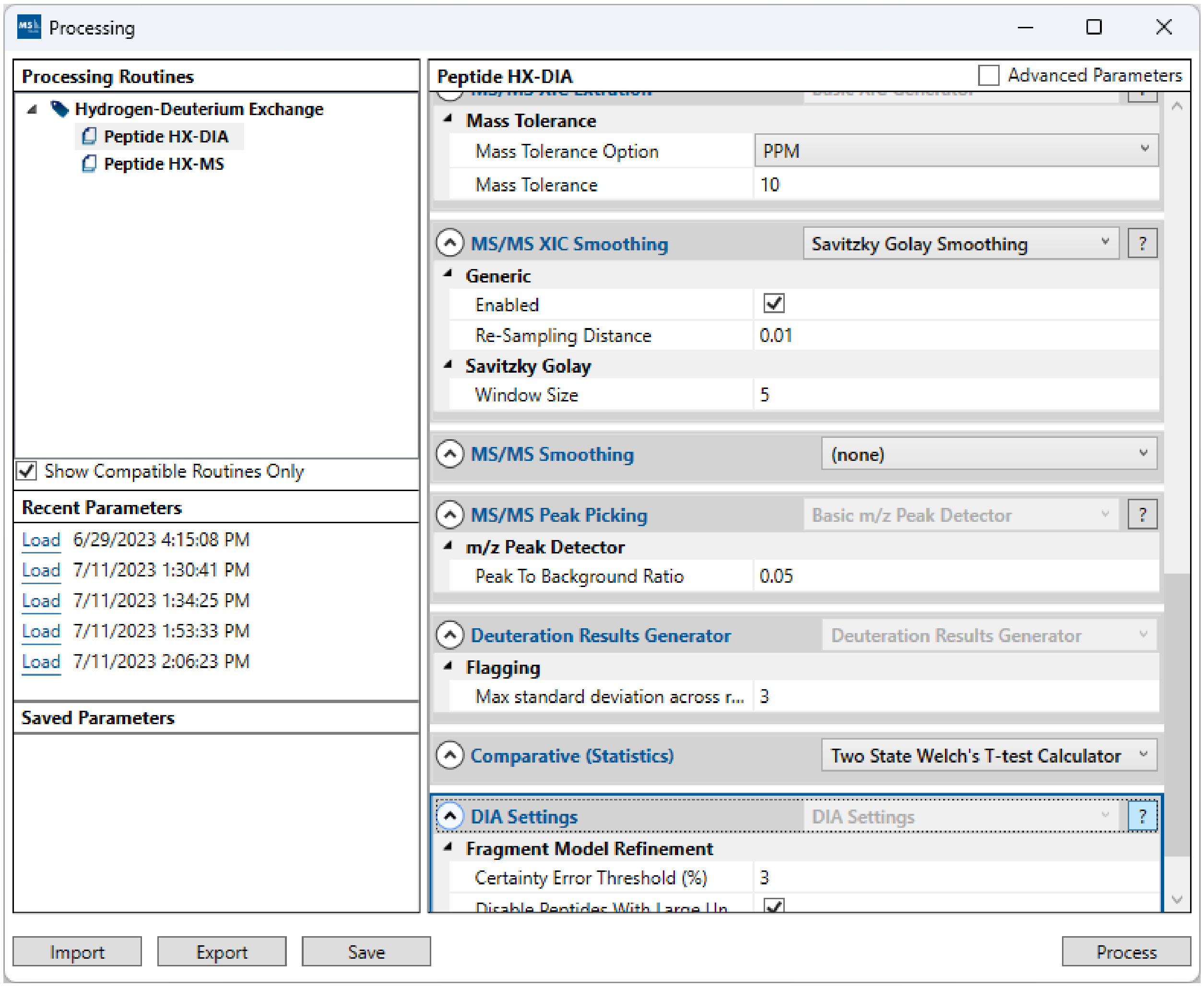


Once everything is set correctly, click “Process”. The total processing time can span from minutes to hours, depending on the number of runs and the number of peptides.

#### Results

After processing is finished, the result will be saved to disk inside the “Results” folder of the project and will be auto-selected in the left-hand-side project tree. AutoHX calculates the “Rescued” deuteration in addition to the standard MS1-based deuteration using the MS2 DIA fragments. The “Final” deuteration for each peptide is based on a decision whether to use the MS1 or MS2 (rescued) deuteration, depending on the quality of the MS1 and MS2 data. For the most part, MS2 deuteration is used for the “Final” deuteration because of its measurement redundancy -- multiple fragments representing the same peptide deuteration during full scrambling, even if the precursor is overlapped in MS1. All of the deuteration values for each peptide are displayed in the right-hand-side Properties tab (per replicate).

The “Manual Validation Tab” in AutoHX is mainly used to inspect the results. We generally don’t recommend any manual intervention after the results are generated because peptides have already gone through multiple stages of strict filtering. To view the decisions for the full set of peptides, including ones which don't pass the final filters, you can disable the hide fragged peptides options in the Processed Peptides list (Major and Critical).

The visualization from HX-DEAL have been updated to show the final AutoHX results. Most notably, the Woods plot now displays certainty bars for each peptide which are determined by the degree of consensus on the peptide deuteration after sampling different sets of fragments. Additionally, the Sequence Coverage view provides a way to map the peptide deuteration or differences onto a 3D structure via the "Export Pymol" button. This button generated a script that can be loaded in PyMOL to colorize the 3D structure based on the data displayed in the Sequence Coverage view.

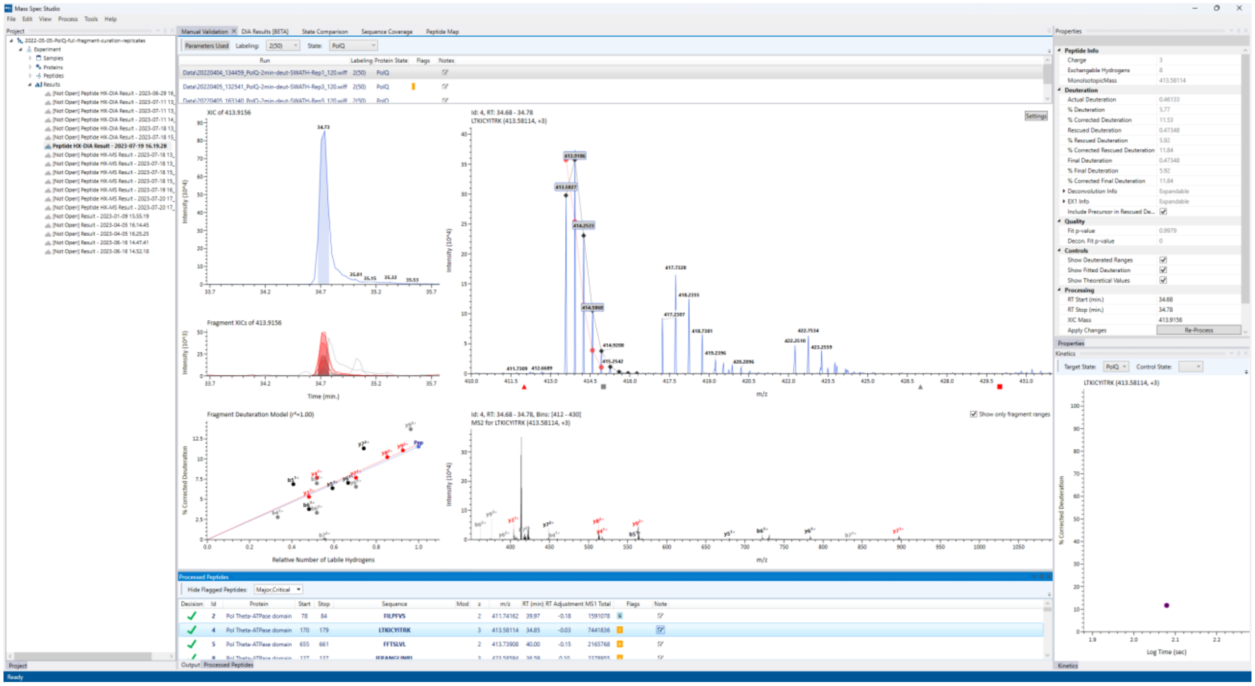

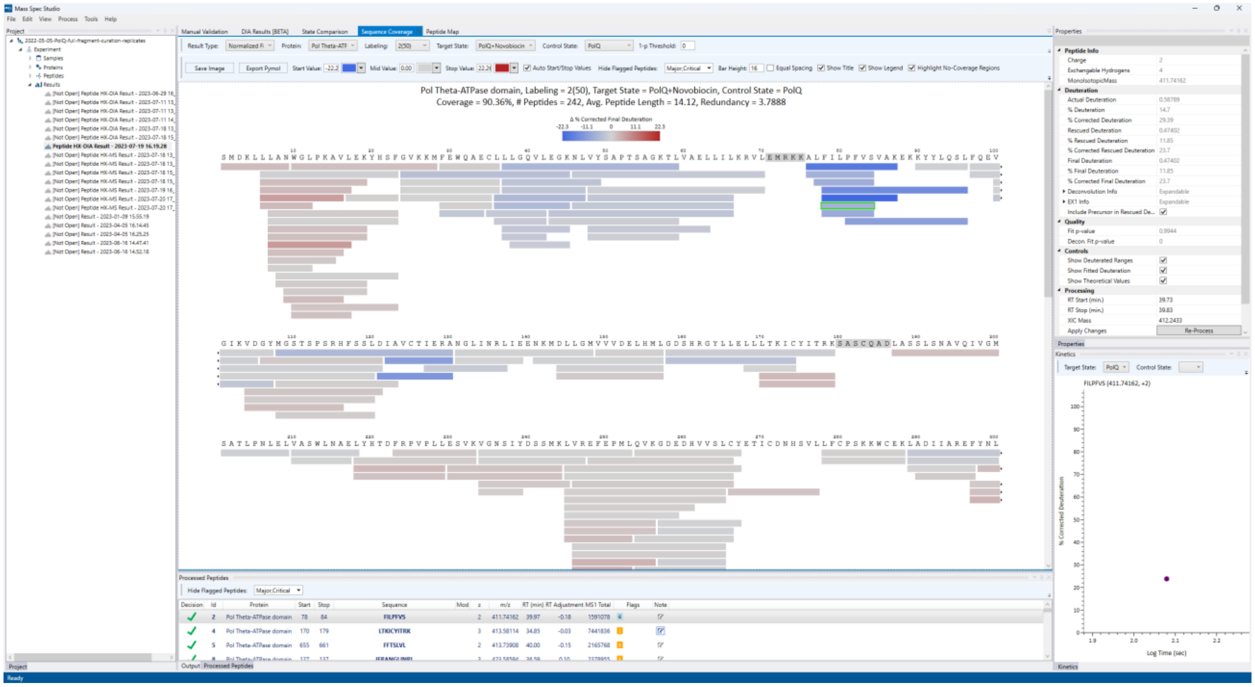
Mass Spec Studio
Manual Validation
DIA Results [BETA]
State Comparison
Sequence Coverage
Peptide Map
Pol Theta-ATPase domain, Labeling = 2(50), Target State = PolQ+Novobiocin, Control State = PolQ
Coverage = 90.36%, # Peptides = 242, Avg. Peptide Length = 14.12, Redundancy = 3.7788
Processed Peptides
Properties
Kinetics
FILPFVS (411.74162, +2)
Log Time (sec)

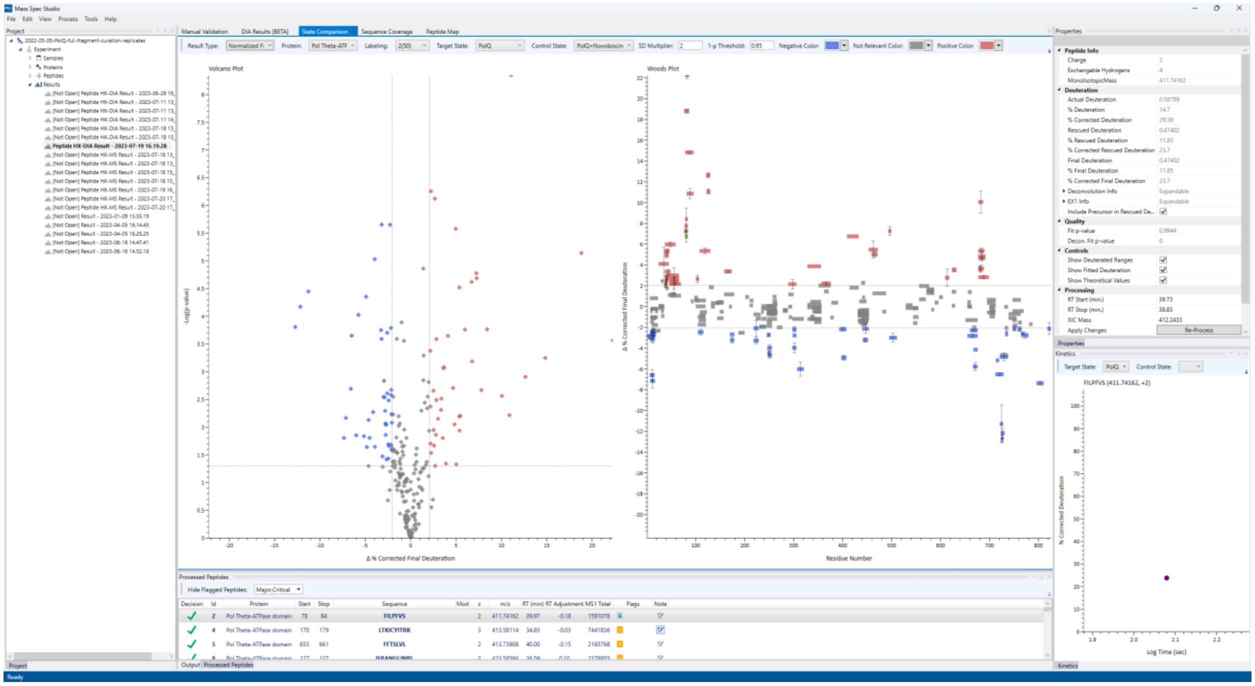
Mass Spec Studio
Manual Validation
DIA Results [BETA]
State Comparison
Sequence Coverage
Peptide Map
Volcano Plot
Woods Plot
Δ % Corrected Final Deuteration
Residue Number
Processed Peptides
Properties
Kinetics
FILPFVS (411.74162, +2)
Log Time (sec)

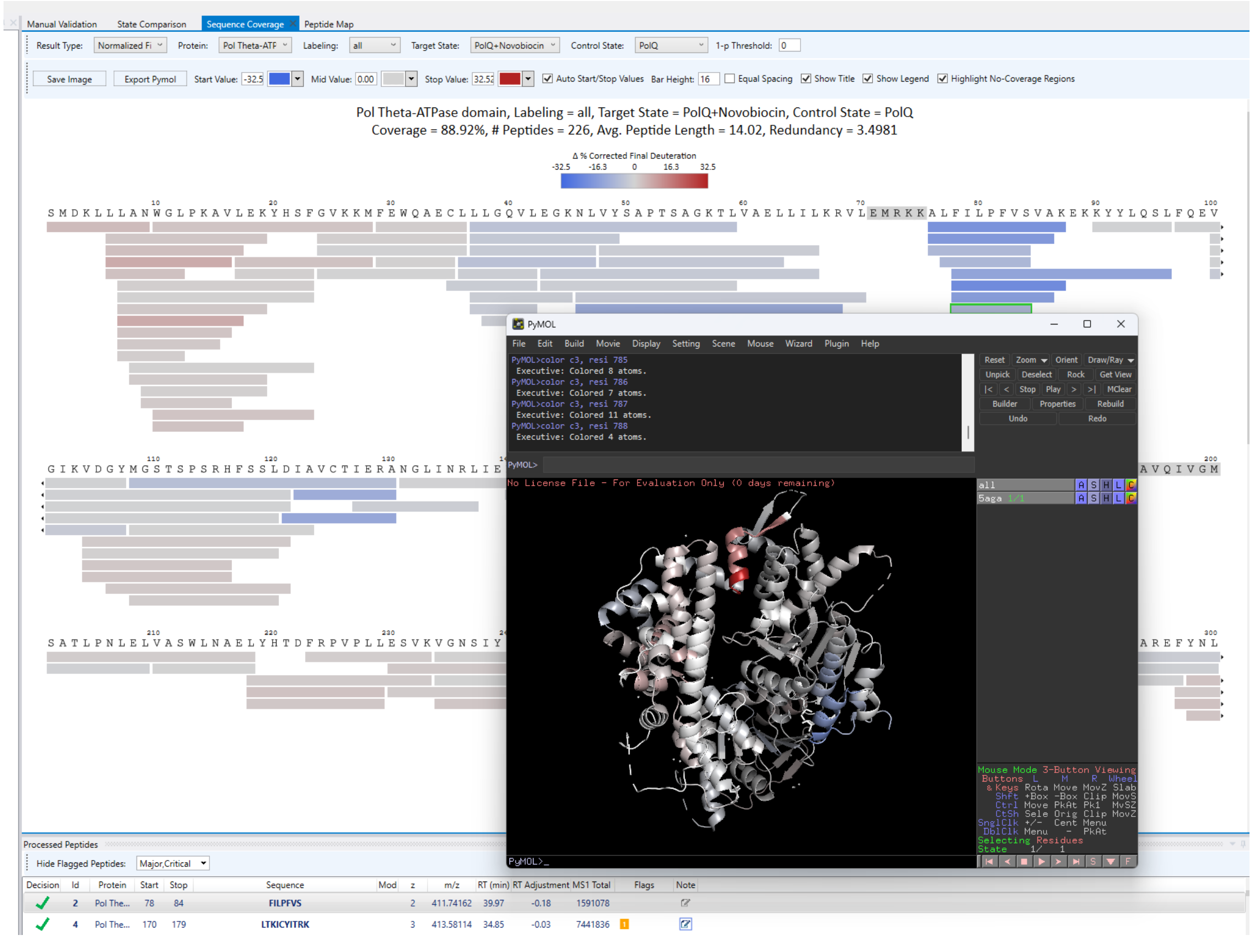

### Export

To export the results, click on File -> Export and follow the export wizard. This follows the standard HX-DEAL export wizard which allows you to select and configure everything from peptide-level .csv exports to snapshots of the commonly used visualizations (Sequence Coverage, Peptide Map, Kinetics, Woods Plot, Volcano Plot). Note that if you wish to export the visualizations, they will need to be configured once before the export. The "Configuration" column will display all the selected exports that require manual configuration before the final export bundle is created. Configuration usually involves setting a template for how the visual should look in the final export bundle (size, DPI, fonts, etc.).

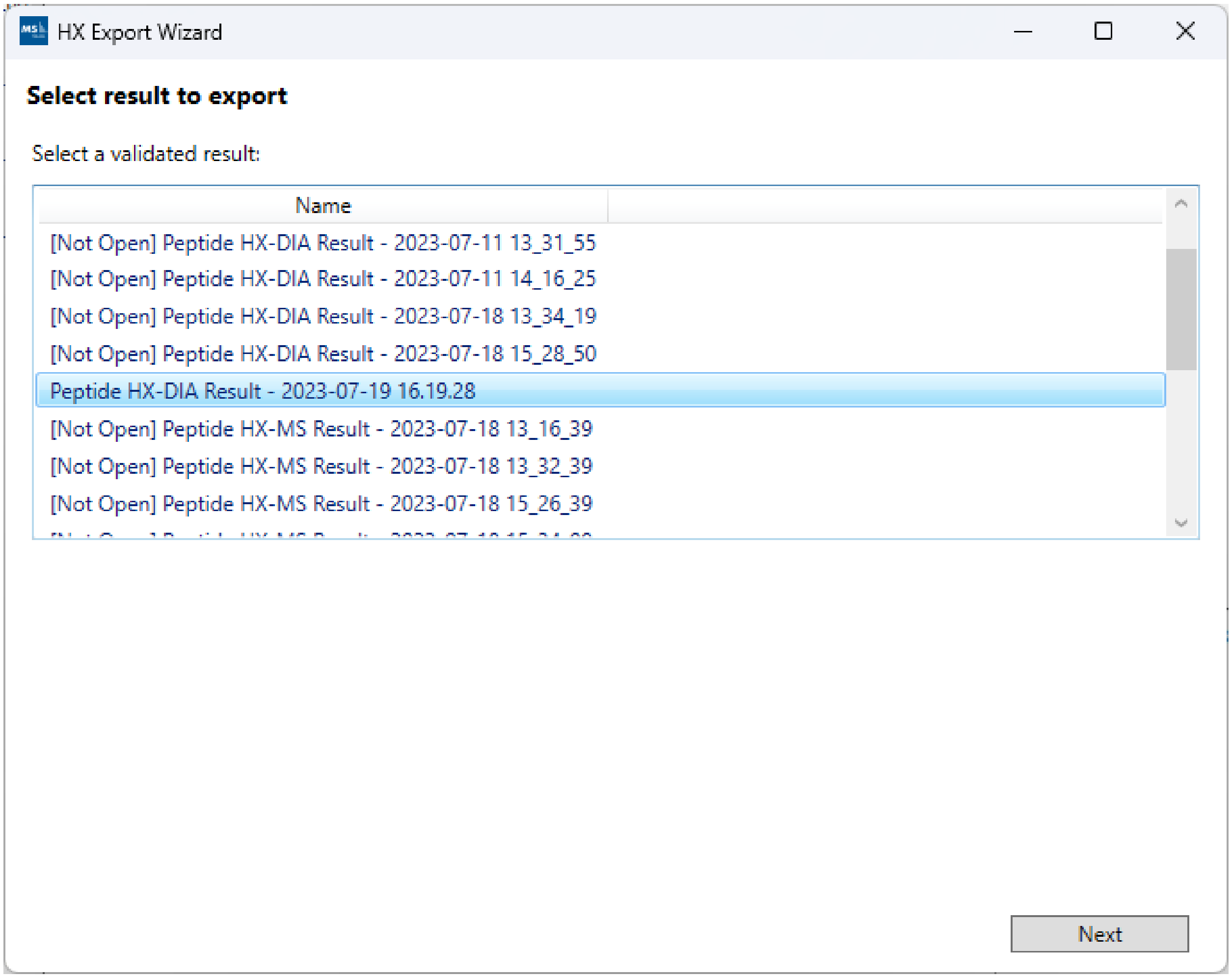
HX Export Wizard
Select result to export
Select a validated result:
Name
[Not Open] Peptide HX-DIA Result - 2023-07-11 13_31_55
[Not Open] Peptide HX-DIA Result - 2023-07-11 14_16_25
[Not Open] Peptide HX-DIA Result - 2023-07-18 13_34_19
[Not Open] Peptide HX-DIA Result - 2023-07-18 15_28_50
Peptide HX-DIA Result - 2023-07-19 16.19.28
[Not Open] Peptide HX-MS Result - 2023-07-18 13_16_39
[Not Open] Peptide HX-MS Result - 2023-07-18 13_32_39
[Not Open] Peptide HX-MS Result - 2023-07-18 15_26_39
Next

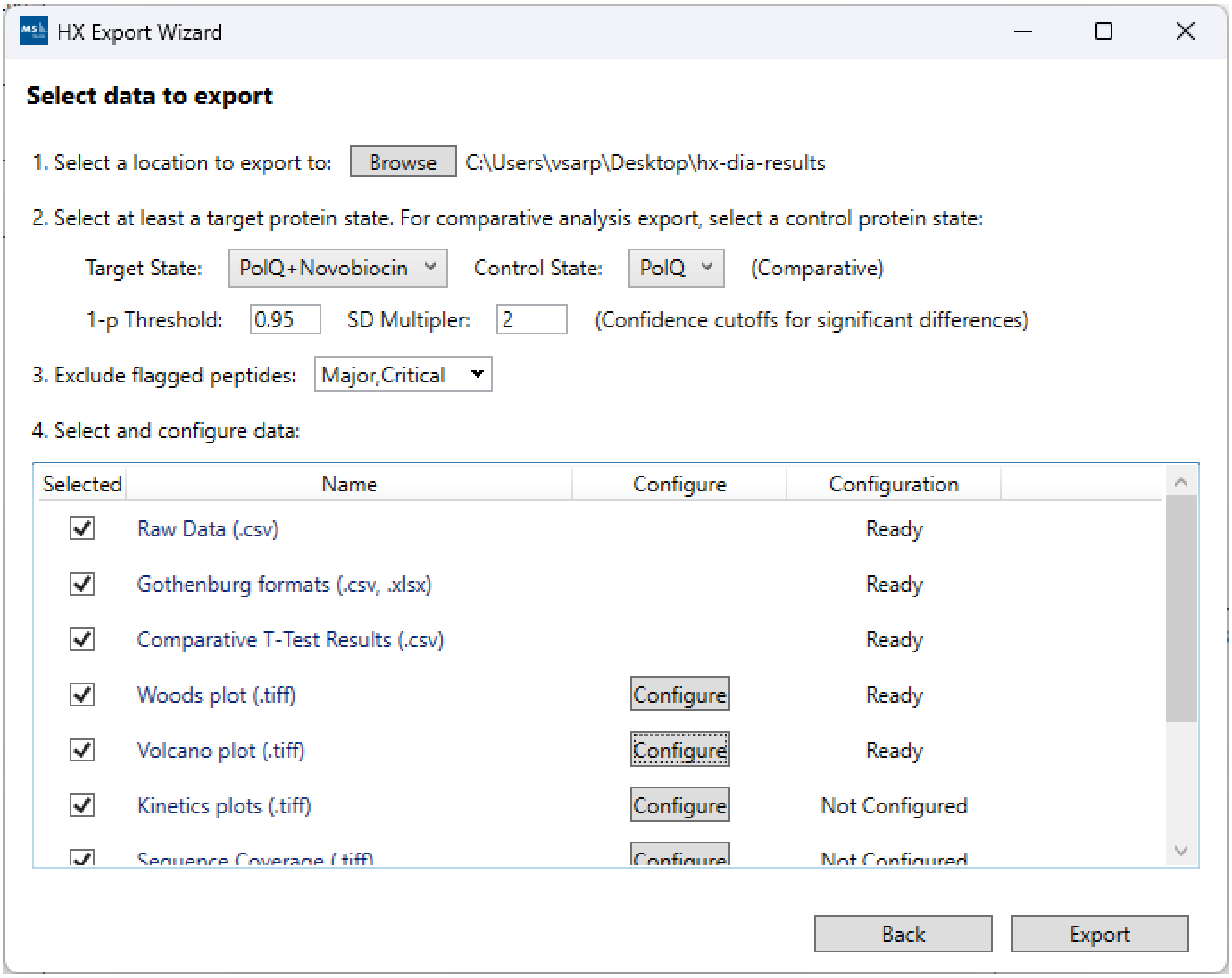
HX Export Wizard
Select data to export
1. Select a location to export to:
Browse
C:\Users\vsarp\Desktop\hx-dia-results
2. Select at least a target protein state. For comparative analysis export, select a control protein state:
Target State:
PolQ+Novobiocin
Control State:
PolQ
(Comparative)
1-p Threshold:
0.95
SD Multipler:
2
(Confidence cutoffs for significant differences)
3. Exclude flagged peptides:
Major,Critical
4. Select and configure data:
Selected
Name
Configure
Configuration
Raw Data (.csv)
Ready
Gothenburg formats (.csv, .xlsx)
Ready
Comparative T-Test Results (.csv)
Ready
Woods plot (.tiff)
Configure
Ready
Volcano plot (.tiff)
Configure
Ready
Kinetics plots (.tiff)
Configure
Not Configured
Back
Export
